# Relationship between neural phase-locked responses to speech and perception of speech in noise in young and older adults

**DOI:** 10.1101/794313

**Authors:** Guangting Mai, Peter Howell

## Abstract

Auditory phase-locked responses are affected by aging and it has been proposed that this increases the challenges experienced during speech perception in noise (SPiN). However, this proposal lacks direct support. This issue was addressed by measuring speech-evoked phase-locked responses at subcortical (frequency-following responses, FFRs) and cortical (theta-band phase-locking, θ-PLV) levels, and studying the relationship between these phase-locked responses and SPiN (word report accuracies of sentences listened to in noise) in adults across a wide age-range (19-75 years old). It was found that: (1) FFR magnitudes declined with age after hearing loss was controlled for; (2) θ-PLV increased with age, suggesting cortical hyperexcitability in audition; (3) SPiN correlated positively with FFR magnitudes obtained in quiet and with θ-PLV obtained in noise, suggesting that impacts of aging (smaller FFR magnitudes and greater θ-PLV) on SPiN differ at subcortical and cortical levels. Thus, the current study provided evidence for different mechanisms at subcortical and cortical levels through which age affects speech-evoked phase-locked activities and SPiN.

## 1 Introduction

Older adults frequently experience increased difficulty with speech perception in noise (SPiN) (Hume and Dubno, 2010). The factors governing this change with age have been investigated extensively. Declines in cognitive functions such as working memory and attention (Lin et al., 2013) as well as hearing loss, degrade SPiN. Older adults, for example, are poorer than young adults at attending to auditory targets when they have to ignore distractor sounds (Andres et al., 2006). This reduction in attentional ability could lead to poor processing of target speech sounds in noisy environments (Tun et al., 2002, 2009). In addition to degraded cognitive functions, older adults have reduced spectro-temporal acuity that could affect SPiN. This is evident in reduced frequency selectivity (Sommers and Gehr, 1998), impaired temporal gap detection (Schneider and Hamstra, 1999; Humes et al., 2009; Harris et al., 2010) and poorer temporal fine structure (TFS) sensitivity (Hopkins and Moore, 2011; Fullgrabe et al., 2015).

The present study focused on the recent claim that impaired SPiN in older adults is due to degraded temporal encoding of speech sounds (Anderson et al., 2011, 2012; Presacco et al., 2016a). The speech-evoked frequency following responses (FFRs) that originate primarily from the auditory brainstem (Chandrasekaran and Kraus, 2010; Bidelman 2018) require precise temporal processing. FFRs reflect the neural phase-locked activity in response to speech envelope modulations at the fundamental frequency (F_0_) (FFR_ENV_F0_) and TFS for higher harmonics in speech (FFR_TFS_) (Aiken and Picton, 2008; Skoe and Kraus, 2010). Previous studies have shown that FFRs are smaller in magnitude in older than in young adults (Anderson et al., 2012; Presacco et al., 2016a) which may indicate that reduced FFR magnitudes lead to impaired SPiN in older adults (Anderson et al., 2012). Furthermore, FFRs are associated with SPiN in older adults (Anderson et al., 2011; Fujihira and Shiraishi, 2015; Mai et al., 2018). Specifically, greater FFR_ENV_F0_ magnitude was associated with better SPiN with speech-shaped noise (Anderson et al., 2011). Greater magnitude of FFR_TFS_ in the resolved harmonics region has also been associated with better SPiN when there is reverberation (Fujihira and Shiraishi, 2015) and multi-talker babble noise (Mai et al., 2018).

Cortical phase-locked activity during speech processing also requires neuro-temporal precision. Auditory phase-locking at low frequencies, especially in the theta range (4 ∼ 8 Hz), tracks slowly-fluctuating speech envelopes at the cortical level (Luo and Poeppel, 2007; Howard and Poeppel, 2010; Peelle et al., 2013). Theta phase-locking (θ-PLV) in response to amplitude-modulated tones, increases with age (Tlumak et al., 2015; Goossens et al., 2016). This is consistent with prior findings that showed older adults had increased auditory-evoked responses (Alain et al., 2014; Herrmann et al., 2013, 2016) and that cortical tracking of speech envelopes was enhanced (Presacco et al., 2016a). Previous studies showed that theta phase-locking to auditory stimuli with envelope modulations at similar frequencies can predict greater neural firing (Ng et al., 2013) and haemodynamic responses in the auditory cortex (Oya et al., 2018). This indicates that increases in cortical phase-locking could be due to hyperexcitability of the auditory cortex in older adults (Caspary et al., 2008). The increased cortical excitability may also alter the balance between inhibitory and excitatory neural processes in older adults that changes network connectivity and over-represents speech envelopes relative to other speech features (Presacco et al., 2016a).

Despite the reported age effects on subcortical and cortical auditory phase-locked responses, it is unclear at present whether they are related to poor SPiN due to aging. Although subcortical speech-evoked FFRs are associated with SPiN in older adults (Anderson et al., 2011; Fujihira and Shiraishi, 2015; Mai et al., 2018), there is no definitive evidence that shows how age effects on FFRs are related to impaired SPiN. For instance, two recent studies (Presacco et al. 2016a; Schoof and Rosen, 2016) tested the relationship between FFR magnitudes and SPiN in adults using young (< 30 years old) and older (> 60 years old) age groups. Although declines in both FFRs and SPiN were found in older adults compared to young adults, neither study found significant FFR-SPiN correlations. The failure to find neural-behavioral correlations at subcortical levels in Schoof and Rosen (2016) and Presacco et al. (2016a) could be because they were conducted in older adults with relatively normal audiometric hearing (thresholds < 30 dB HL at frequencies ≤ 4 kHz). Hence, there were no participants with significant hearing decline due to normal aging, meaning that the older test group did not represent the complete range of hearing present in typical aging populations (Gopinath et al., 2009; Humes et al., 2010).

At the cortical level, Presacco et al. (2016a) argued that increased phase-locking to speech in older adults may reflect a loss of excitation-inhibition balance. It is not clear whether such loss is a mechanism for impaired SPiN during aging and Presacco et al. (2016a) failed to find any correlation between cortical neural activity and SPiN. This absence of correlation may be because different types of background noises were used when neural data and SPiN were measured (single-talker background during neural recording and four-talker babble noise in SPiN tasks; see Presacco et al. (2016a)). On the other hand, studies have shown that greater cortical phase-locking to speech is associated with higher intelligibility of degraded speech stimuli in normal-hearing young adults (Ahissar et al., 2001; Peelle et al., 2013; Doelling et al., 2014) as well as better SPiN in older adults (Mai et al., 2018). These findings are consistent with other studies which showed that greater magnitudes of cortical auditory evoked potentials (CAEPs) can predict better SPiN in both young and older adults (Billings et al., 2013, 2015). Therefore, the relation between age effects on cortical processing and SPiN has not been clarified so far, i.e., whether increased cortical phase-locking according to aging relates to impaired SPiN (due to dampened excitation-inhibition balance) or better SPiN (due to greater sensitivity to speech).

The present study addressed whether age effects on subcortical/cortical phase-locked encoding of speech were associated with impaired SPiN. Behavioral and neural assessments were conducted in healthy adults across a wide age-range (19-75 years). Older adults in the present study had audiometric thresholds at frequencies between 2 and 4 kHz indicative of normal hearing to mild/moderate hearing loss. Therefore individual variability associated with peripheral hearing that occur during normal aging was present in the sample (Gopinath et al., 2009; Humes et al., 2010). For the behavioral assessment, participants completed SPiN tasks under two types of background noise: steady-state speech-shaped noise (SpN) and 16-talker babble noise (BbN). For the neural assessments, participants listened to a repeated syllable under the same types of noise as in the behavioral assessment, whilst speech-evoked phase-locked activity was recorded at both subcortical (FFRs) and cortical (θ-PLV) levels using scalp-electroencephalography (EEG). SPiN and the neural signatures were compared across the two age groups and multiple linear regressions were conducted to investigate whether the age-related neural signatures were associated statistically with SPiN.

Based on past evidence, it was predicted that older, relative to young, adults would have: (1) smaller subcortical (FFRs) magnitudes (Anderson et al., 2012; Presacco et al., 2016a); (2) greater cortical (θ-PLV) phase-locked responses to speech (Presacco et al. 2016a); and (3) decreased SPiN (Hume and Dubno, 2010). The predictions for testing the hypotheses that age effects on neural measures relate to behavioral performance are that decreased SPiN with age should be statistically associated with: (1) reduced FFR magnitudes; and (2) greater θ-PLV. At the same time, this study explores which neural (subcortical and/or cortical) signatures optimally model SPiN which is an issue that is not clear to date.

## 2 Methods

The present study followed the same procedure and used parts of the older adults’ data from Mai et al. (2018)^1^.

### 2.1 Participants

Participants comprised 23 young (19-42 years; Mean ± SD = 26.3 ± 5.5 years; 15 males) and 18 older adults (53-75 years; Mean ± SD = 67.0 ± 5.6 years; 7 males). All were native UK English speakers with no reports of neurological diseases, language-related or psychiatric problems. **Figure 1** (left panel) shows the pure-tone audiometric thresholds (PTAs) for frequencies 0.25 ∼ 8 kHz (averaged across both ears) measured using an MA41 Audiometer (MAICO Diagnostics, Germany). All young participants had normal hearing (PTA ≤ 25 dB HL). In older participants, inter-individual variability was high particularly at frequencies of 2 kHz and above. The older adults showed significantly higher low-frequency PTAs (PTA_Low_; averaged across 0.25 ∼ 1 kHz) and high-frequency PTAs (PTA_High_; averaged across 2 ∼ 4 kHz)^2^ compared to the young group (both *p*s < 10^−6^). The boxplot (**Figure 1**, right panel) indicated that PTA_High_ in the older group ranged from normal hearing (≤ 25 dB HL) to mild/moderate (25 ∼ 50 dB HL) hearing loss, comparable with the distribution pattern reported in other older samples (Gopinath et al., 2009; Humes et al., 2010). A two-way mixed-design ANOVA was conducted for PTA with factors of Frequency (PTA_High_ vs. PTA_Low_) and Age Group (young vs. older). A significant [Frequency × Age Group] interaction occurred (*p* = 0.001), indicating that older adults had significantly greater declines in hearing at the high compared to the low frequencies. Since PTAs differed across frequencies and age groups, PTA_Low_ and PTA_High_ were used as well as PTA averaged across the wider frequency range (0.25 ∼ 4 kHz; PTA_Wide_) as separate covariates and predictors during statistical analyses (see *2.4*).

**Figure 1.**
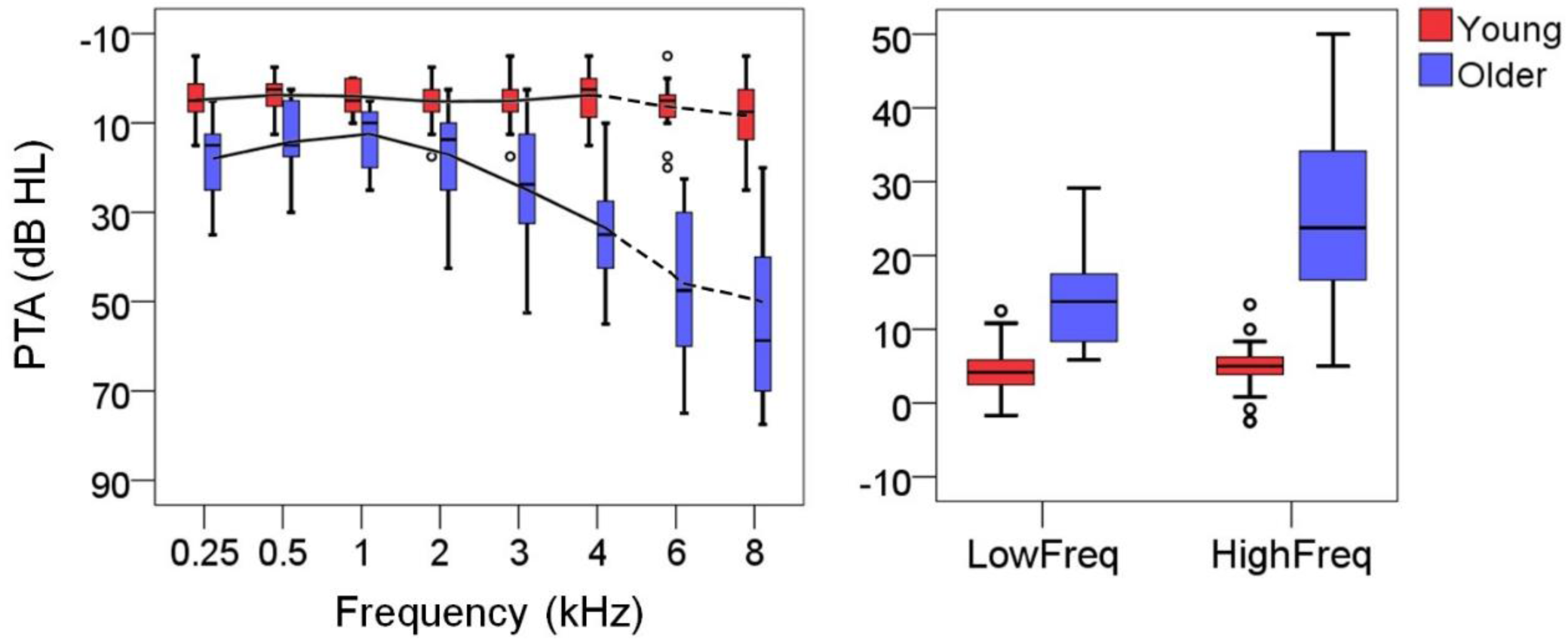
Boxplots of pure-tone audiometric thresholds (PTAs) for the young (red) and older (blue) adults averaged across ears. The left panel shows the audiogram for the range from 0.25 to 8 kHz^3^. Data for 6 and 8 kHz (dashed lines) were not used in the subsequent statistics because spectral distribution of the speech stimuli used in the present study only extended to 4 kHz. The right panel shows the PTAs at the low (0.25 ∼ 1 kHz) and high frequency (2 ∼ 4 kHz) ranges.

### 2.2 Behavioral experiment

SPiN tasks involved participants listening to BKB sentences (Bench et al., 1979) under two types of background noise: steady-state speech-shaped (SpN) and 16-talker babble (BbN) noise. All sentences were pre-recorded utterances spoken by a male British English speaker whose F_0_ ranged from 80 to 200 Hz. Each sentence included three key (content) words, e.g., “The clown has a funny face” with key words “clown”, “funny” and “face”. BbN was a mixture of 16 different utterances spoken by 16 male British English speakers with similar voice quality to the target speaker. SpN was formed by randomizing the phases of the long-term spectrum of BbN and transforming the spectrum back to the time domain. As a result, SpN has the same long-term power spectrum as BbN and stable time-domain properties (Rosen et al., 2013).

Participants were seated comfortably in a sound-treated booth facing a Fostex 6301B loudspeaker (Canford Group Ltd.) at zero-degree azimuth. Distance between the loudspeaker and participants’ ears was constant at 1 meter. After eight trials of practice, participants listened to two different sets of 30 sentences (for the backgrounds of SpN and BbN, respectively) at an intensity at this distance of 70dB SPL. Participants repeated as many words as they could from each sentence. Sentences were presented via Matlab 2010a (Mathwork, USA) and SNR varied adaptively to track for the speech reception threshold (SRT, Plomp and Mimpen, 1979) at which 50% of words were correct. For each background type, the first sentence was played at a relatively high SNR (8 and 10 dB for SpN and BbN, respectively). SNR was decreased by 4 dB for subsequent sentences until < 50% words correct (i.e., < 2 words) were reported. SNR was then increased/decreased by 2 dB when word correctness was less/more than 50% in each of the following sentences. SRT was calculated by linear interpolation using the two SNRs which had > 50% and < 50% correct across the minimal step distance (i.e., 2 dB).

### 2.3 EEG experiment

#### 2.3.1 Acoustic stimuli

Participants listened to a repeatedly-presented, 120-ms-long /i/ syllable produced by a male speaker (**Figure 2A**). The F_0_ contour of the syllable fell from ∼ 160 to ∼110 Hz (**Figure 2B**). The F_0_ contour covered a similar frequency range and direction of change as those in the F_0_s of the target speaker in the BKB sentences used in the SPiN tasks (BKB sentences are narratives that generally have a falling F_0_ contour). The three formants in the syllable were at ∼ 280 Hz (F1), ∼ 2400 Hz (F2) and ∼ 3100 Hz (F3). The amplitude envelope profile was stable except that 5-ms-long rising and falling cosine windows were applied at the onset and offset to avoid transients.

**Figure 2.**
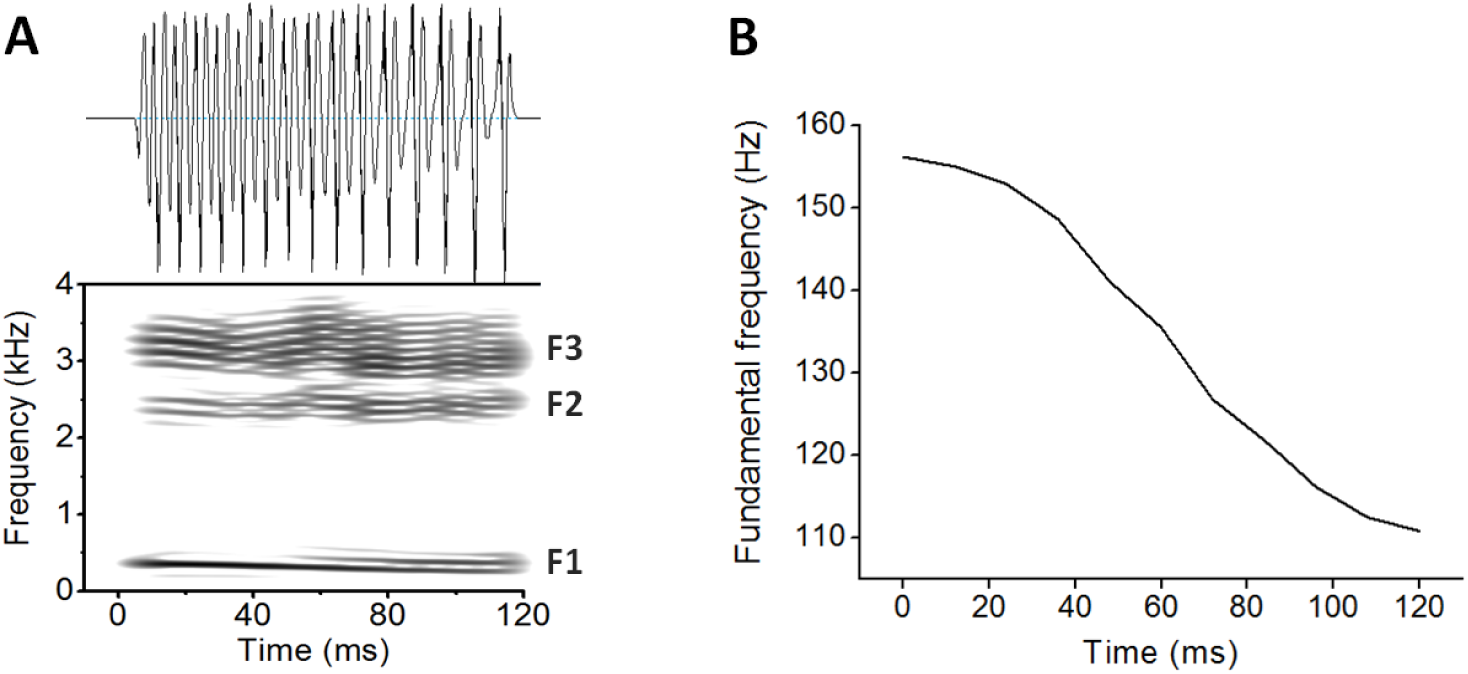
The syllable /i/ used during EEG recording. **(A)** The temporal waveform (top) and spectrogram (bottom) of the syllable. F1, F2 and F3 frequencies are around 280, 2400 and 3100 Hz, respectively. **(B)** The falling F_0_ contour ranging from around 160 to 110 Hz obtained by autocorrelation. The waveform, spectrogram and F_0_ were generated via PRAAT (Boersma and Weenink, 2013).

The syllable was presented repeatedly at both original (positive) and inverted (negative) polarities in random order with inter-stimulus intervals (ISIs) that varied randomly between 60 and 120 ms (syllable repetition rate was approximately 5 syllables per second). The stimuli were presented in quiet, SpN and 16-talker BbN backgrounds (the last two were the same backgrounds that were used in the SPiN tasks). The SNRs were set at −1 dB, which led to neural responses that correlated significantly with SPiN in older adults (Mai et al., 2018). There were 6400 sweeps under each background type (3200 sweeps for each polarity). Recordings at each background type were split into 16 segments of equal duration giving 48 segments in total with 400 sweeps per segment. The segments were played in succession in an intermixed order.

#### 2.3.2 EEG data acquisition

Scalp-EEGs were recorded on an ActiveTwo system (Biosemi, The Netherlands) at a sampling rate of 16384 Hz. Three active electrodes were placed at Cz (vertex), C3 and C4 according to the 10/20 configuration. Cz was used to obtain FFRs (Skoe and Kraus, 2010). Cortical responses were measured on C3 and C4 that reflects activity in the auditory cortex (Carpenter and Shahin, 2013; Noguchi et al., 2015) and allows reliable cortical phase-locked activity that is significantly associated with SPiN to be recorded (Mai et al., 2018). Bilateral earlobes were used as the reference. Ground electrodes were CMS/DRL. Electrode impedance was kept below 35 mV. The experiment was conducted in an electromagnetic-shielded and sound-treated booth. The stimuli were played via a Rogers LS3/5A loudspeaker (Falcon Acoustics, UK) at zero-degree horizontal azimuth relative to participants’ heads when they were reclined (the chair was adjustable). The stimulus level (measured across time including ISIs) at the distance between the loudspeaker and participants’ ears (constant at 1 meter) was calibrated at 74.5 dB before background noise was added. The stimulus level was at 79.5 dB after either SpN or BbN was added.

Participants were instructed to relax, close their eyes and keep still in order to avoid movement artefacts. They did not have to make any response to the stimuli (passive listening) and they were not stopped from falling asleep. A webcam monitored the participants throughout the test and no significant changes in head or body position were observed. Participants were not stopped from falling asleep because another purpose of the current experiment was to study the effects of arousal on speech-evoked neural processing across ages (Mai et al., 2019). This investigation was separate from the present paper. As Mai et al. (2019) found that arousal significantly affected the phase-locked responses, only EEG data from periods with high arousal were used here (see *2.4.3* for details).

### 2.4 Signal processing for EEG data

The signal processing procedure used Matlab 2014a (Mathwork, USA).

#### 2.4.1 Frequency following responses (FFRs)

EEGs at Cz were re-referenced to the average of bilateral earlobes and bandpass filtered between 70 and 4000 Hz using a zero-phase 2nd-order Butterworth filter. Baseline was adjusted using the pre-stimulus period of 50 ms. Sweeps exceeding ± 25 μV were rejected to exclude movement artefacts. FFRs with positive (FFR_pos_) and negative (FFR_neg_) polarities were obtained by averaging across sweeps with their respective polarities. FFRs that represent envelope modulations (FFR_ENV_) and TFS (FFR_TFS_) respectively were obtained by addition and subtraction of FFR_pos_ and FFR_neg_ that were then divided by 2 (Aiken and Picton, 2008). **Figure 3** shows an example of FFRs obtained in the present study (FFRs of a single participant recorded in BbN).

**Figure 3.**
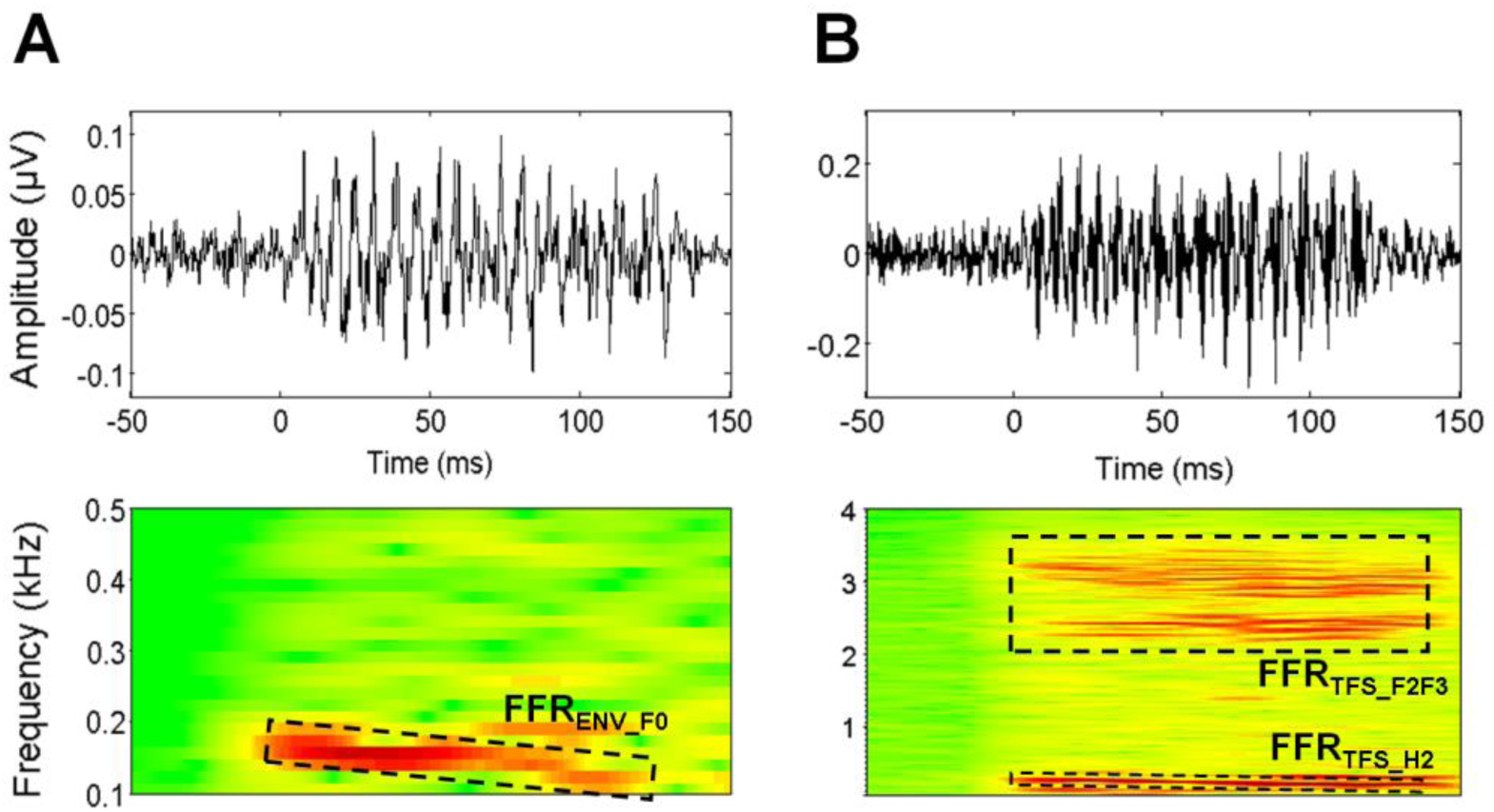
FFR waveforms (top) and spectrograms (bottom) of a single participant recorded in BbN. **(A)** FFR_ENV_ (bandpass filtered at 70 ∼ 2000 Hz); **(B)** FFR_TFS_ (bandpass filtered at 70 ∼ 4000 Hz). The waveforms were based on sweeps with normalized numbers (1450 ∼ 1550) during the high arousal state (see *2.4.3 and 2.4.4*). FFR_ENV_F0_ (at F_0_ range between 160 and 110 Hz), FFR_TFS_H2_ (at H2 range between 220 and 320 Hz) and FFR_TFS_F2F3_ (at F2-F3 range between 2000 and 4000 Hz) are indicated by the boxes surrounded by dashes and their labels. ‘0’ corresponds to the syllable onset.

Three FFR magnitudes were measured: (1) FFR_ENV_F0_ that represents neural encoding of envelope modulations at F_0_, quantified as the magnitude along the F_0_ trajectory using FFR_ENV_ (**Figure 3A**); FFR_TFS_H2_ that represents neural encoding of TFS at the resolved harmonics region (2^nd^ harmonics H2 at 220 ∼ 330 Hz in the neighborhood of F1), quantified as the magnitudes along the H2 trajectory using FFR_TFS_ (**Figure 3B**); and (3) FFR_TFS_F2F3_ that represents neural encoding of TFS in the unresolved harmonics region (frequency range around F2 and F3), quantified as the magnitudes along the F2 and F3 trajectories using FFR_TFS_ (**Figure 3B**). The procedures for spectral magnitude calculations followed Mai et al. (2018).

First, ENV-F_0_ (F_0_ based on the acoustic envelope), H2, F2 and F3 trajectories of the /i/ syllable were calculated. To obtain the ENV-F_0_ trajectory, a set of 40-ms sliding windows (1-ms per step) was applied to the syllable’s Hilbert envelope. Each 40-ms segment was Hanning-windowed, zero-padded to 1 second (to achieve 1 Hz frequency resolution) and Fourier-transformed. The frequency with the highest Fourier magnitude between 110 and 160 Hz (the F_0_ range) was chosen as the F_0_ value at each step. H2, F2 and F3 trajectories were obtained in the same way, except that: 1) sliding windows were applied to the syllable rather than the Hilbert envelope; 2) H2 values were selected within the H2 range (220 ∼ 320 Hz); 3) instead of choosing values based on the Fourier spectrum after zero-padding, F2 and F3 values were chosen based on the spectral profile via cepstral smoothing (Proakis and Manolakis, 2007) in the F2 and F3 ranges respectively (2200 ∼ 2600 Hz and 2800 ∼ 3500 Hz). Second, to calculate the FFR_ENV_F0_ magnitude, the same set of 40-ms sliding windows was applied to FFR_ENV_ with Hanning-windowing, zero-padding and Fourier-transforms. The mean log-magnitude was measured across a 20 Hz bandwidth centered at the frequency of the ENV-F_0_ trajectory at that step. The magnitudes were then averaged across all steps along the ENV-F_0_ trajectory. FFR_TFS_H2_ and FFR_TFS_F2F3_ magnitudes were obtained in the same way, except that: 1) the procedure was applied on FFR_TFS_ along the H2 (for FFR_TFS_H2_) and the F2 and F3 (for FFR_TFS_F2F3_) trajectories; 2) instead of obtaining magnitudes based on the Fourier spectrum after zero-padding, FFR_TFS_F2F3_ magnitude at each step was the summed magnitude of the spectral profile (via cepstral smoothing) across a 150 Hz and 300 Hz bandwidth respectively centered at F2 and F3 of the syllable at that step.

In addition, neural transmission from the cochlea to the auditory brainstem for FFR_ENV_ takes between 5 and 10 ms (Chandrasekaran and Kraus, 2010; Skoe and Kraus, 2010), while FFR_TFS_ occurs at earlier stages in the auditory periphery (Aiken and Picton, 2008). Hence, the maximum magnitude for time lags in the range 8 to 13 ms and 3 to 8 ms (at 1-ms steps; including an additional 3 ms of air transmission from the loudspeaker to the cochlea) were used as the final FFR_ENV_F0_ and FFR_TFS_ (FFR_TFS_H2_ and FFR_TFS_F2F3_) magnitudes, respectively.

As well as FFR magnitudes, inter-trial phase-locking values (PLV) at the F_0_ (FFR_PLV_F0_) were also calculated. This was because FFR_PLV_F0_ reflects pure phase-locking that excludes the influence of single-trial spectral magnitudes and has a better signal-to-noise ratio than does FFR magnitudes (Zhu et al., 2013). FFR_PLV_F0_ was calculated in a similar way to FFR_ENV_F0_ along the ENV-F_0_ trajectory, except that, after zero-padding in each step (without Hanning-windowing), PLV was calculated (Morillon et al., 2012) instead of spectral magnitudes:

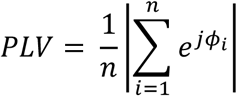

where *n* denotes the total number of sweeps, *ϕ*_*i*_ denotes the Fourier phase value at the frequency of the ENV-F_0_ trajectory for the *i*th sweep at that step, and *j* is 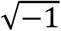. As PLV is restricted to values between 0 and 1, FFR_PLV_F0_ at each step was quantified by logit-transforming PLV to [-∞, +∞], making it appropriate for linear regression analysis (Waschke et al., 2017):

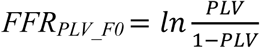

FFR_PLV_F0_ values were then averaged across all steps along the ENV-F_0_ trajectory. The final FFR_PLV_F0_ value was taken as the maximal value for the time lags between 8 and 13 ms as in measurement of the FFR_ENV_F0_ magnitude.

#### 2.4.2 Cortical responses

Cortical responses were measured as theta-band (4 ∼ 6 Hz, to correspond to the stimulus repetition rate of 5 syllables per second) phase-locking values (θ-PLV) at C3 and C4. EEGs were decimated to 1024 Hz, re-referenced to the average of the bilateral earlobes and bandpass filtered (4 ∼ 6 Hz) using a 2nd-order zero-phase Butterworth filter. Sweeps exceeding ± 15 μV on either electrode were rejected (Mai et al., 2018)^4^. θ-PLV time series (*PLV(t)*) were calculated and then logit-transformed:

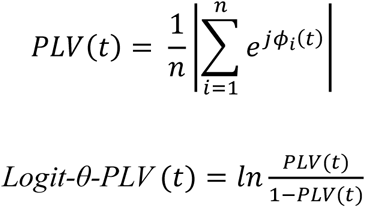

where *n* denotes the total number of sweeps, *ϕ*_*i*_*(t)* denotes the Hilbert phase series of the filtered EEG of the *i*th sweep time-locked to the syllable onset and *j* is 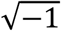. Hilbert phase was used as it reflects phase-locking to stimuli even when EEG amplitude variation occurs due to relaxation and eye closure (Thatcher, 2012). Logit-θ-PLV(t) values were then averaged across the stimulus period (120 ms). Neural transmission from cochlea to auditory cortex takes 10 to 30 ms in primates (Lakatos et al., 2007). Hence Logit-θ-PLV was taken as the maximum value for time lags between 13 and 33 ms (at 1-ms steps) with the 3 ms for air transmission included. Finally, the Logit-θ-PLV(t) was averaged across the two electrodes.

#### 2.4.3 Classification of arousal states

Participants were not required to remain awake during the EEG recording because a separate purpose of the present experiment was to investigate the effect of arousal on speech-evoked responses (Mai et al., 2019). Mai et al. (2019) reported that both subcortical and cortical responses showed significant suppression in low arousal/nREM states compared to high arousal states. This accords with earlier functional imaging work that showed that neural responses to speech in subcortical (Portas et al., 2000) and cortical (Czisch et al., 2004; Wilf et al., 2016) auditory regions reduce during sleep compared to wakefulness. Also, significant correlations between behavioral measures and EEG parameters (FFRs and θ-PLV) were only found in the high arousal state (Mai et al., 2018). Hence, only EEG data from periods with high arousal were used to avoid any influences that arousal has on the neural-behavioral relationship.

Sleep spindles were used to determine arousal state (Martin et al., 2013). Sections of the Cz EEG recordings were categorized into three types of epochs (all epochs were 21-second long) based on the occurrence of sleep spindles: (1) epochs in high arousal states (wakefulness or nREM Stage 1); (2) epochs in low arousal states (nREM Stage 2); and (3) epochs in transition between (1) and (2). After the experiment, participants gave a subjective ranking concerning how much they had slept. There was a significant correlation between the sleep ranking and the percentage of epochs classified as ‘low arousal’ (*p* = 0.002), which validated the spindle-based method. Further methodological details are available in Mai et al (2019).

#### 2.4.4 Normalization of sweep numbers

Robust FFRs require around 1500 artefact-free sweeps (c.f., Dajani et al., 2005; Wong et al., 2007). Hence, participants’ data for a particular background type were not included in subsequent analyses if there were < 1,450 artefact-free sweeps in high arousal epochs for that background type. This resulted in 17%-30% of participants being rejected from further analyses (depending on the types of analyses conducted; see *3.2* and *3.3* for details). Moreover, as magnitudes of phase-locked activity are sensitive to the number of sweeps (Aviyente et al., 2011), problems can arise during statistical analyses if the number of sweeps differ significantly across participants. Therefore, the number of sweeps was normalized to around 1,500 for both FFRs and θ-PLV for each participant in each background type. Normalization was achieved by selecting high arousal epochs at random that included 1,450 to 1,550 sweeps and EEG signatures (magnitudes of FFR_ENV_F0_, FFR_TFS_H2_ and FFR_TFS_F2F3_, and Logit-θ-PLV) were obtained from the selected epochs. The random selection procedure was repeated 100 times, giving 100 estimates for each EEG signature. Averages over the 100 estimates were used in the final statistical analyses. Therefore, this process ensured EEG signatures were based on around 1500 sweeps regardless of artefact rejection rates or different number of epochs of the three arousal states across participants (Mai et al., 2018).

#### 2.4.5 Confirming FFR robustness

FFR magnitudes are small and their robustness was tested by statistically comparing the FFR magnitudes with the EEG noise floors. The noise floors were quantified as the EEG magnitudes at the corresponding frequency range (110 ∼ 160 Hz (F_0_) for FFR_ENV_F0_; 220 ∼ 320 Hz (H2) for FFR_TFS_H2_; 150-Hz bandwidth centred at 2400 Hz (F2) and 300-Hz bandwidth centred at 3100 Hz (F3) for FFR_TFS_F2F3_) at the 50-ms FFR pre-stimulus period (Mai et al., 2018). The quantification procedure was similar to that used in calculating FFR magnitudes, in which a set of 40-ms sliding windows was applied on FFR_ENV_ (for FFR_ENV_F0_) or FFR_TFS_ (for FFR_TFS_H2_ and FFR_TFS_F2F3_) which used 1-ms steps over the pre-stimulus period. Magnitudes of noise floors were measured as the spectral magnitudes (summed magnitude of the cepstrally-smoothed profile for FFR_TFS_F2F3_) across the corresponding frequency ranges averaged across all steps. These were all conducted along with the calculations of FFR magnitudes during the processes for normalization of sweep numbers.

### 2.5 Statistical analyses

Statistical analyses were conducted using SPSS 13.0 (SPSS Inc., USA).

#### 2.5.1 Analyses of variance (ANOVAs) and Analyses of covariance (ANCOVAs)

Mixed-design ANOVAs were conducted with the behavioral (SRT in the SPiN tasks) and EEG signatures (FFR_ENV_F0_, FFR_PLV_F0_, FFR_TFS_H2_, FFR_TFS_F2F3_ and Logit-θ-PLV) as the dependent variables, and with the within-subject factor of Noise Type (SpN and BbN for SRT; Quiet, SpN and BbN for EEG) and the between-subject factor of Age Group (young vs. older) as independent variables. Post-hoc t-tests were conducted if a significant [Noise Type × Age Group] interaction occurred. The ANOVAs were conducted for two reasons: (1) main effects of Age Group were tested to look into the effects of age on SPiN and neural responses to speech; (2) since older adults experience more difficulties in SPiN under BbN compared to other types of noise (Helfer and Freyman, 2008; Schoof and Rosen, 2014), testing the [Noise Type × Age Group] interaction should help reveal what neural mechanisms underlie this.

The EEG signatures were further used in mixed-design ANCOVAs that included PTAs (PTA_Low_, PTA_High_ and PTA_Wide_, averaged across 0.25 ∼ 1 kHz, 2 ∼ 4 kHz and 0.25 ∼ 4 kHz, respectively; all mean-centred) as covariates, with the same within-subject and between-subject factor as in ANOVAs. These tested for age effects after the variability in peripheral hearing loss was controlled for. PTA_Low_ and PTA_High_ reflect hearing loss at low and high frequencies, respectively, while PTA_Wide_ reflects the combined effect of both. The three PTA variables were used as covariates in separate analyses to avoid the risk of collinearity. A concern about the ANCOVAs is that PTAs and Age are correlated (effects of PTA and Age can overlap), hence including PTAs as covariates may carry risks of partially diminishing the Age effects despite controlling for hearing loss. For this reason, ANOVAs and ANCOVAs were both conducted separately. Alpha values for testing significance in ANOVAs/ANCOVAs were not adjusted.

#### 2.5.2 Multiple linear regressions

Multiple linear regressions were then conducted to test for any behavioral-neural relationship using data from both the young and older groups. EEG signatures (FFR_ENV_F0_, FFR_PLV_F0_, FFR_TFS_H2_, and FFR_TFS_F2F3_, and Logit-θ-PLV) were used as predictors of SPiN (SRTs; dependent variables) for the corresponding noise types. Specifically, EEG signatures obtained in SpN were used to predict SRT in SpN, while EEG signatures obtained in BbN were used to predict SRT in BbN. This avoided problems that could arise due to behavioral and neural recordings being made under different types of noise (Presacco et al., 2016a). Additionally, since FFRs in quiet have been suggested to be associated with SPiN (Anderson et al., 2011), we also used EEG signatures in quiet to predict SRTs. Age was not included as a predictor, since the regressions investigated the contributions of age-related factors (which were identified by significant main effects of Age in the ANOVAs), rather than age itself, to SPiN.

The Best-Subset Regression approach was used that selected predictors of EEG signatures that generated the lowest BIC value. This approach provided the optimal model with best goodness of fit and least chance of overfitting (Burnham and Anderson, 2003). PTAs (PTA_Low_, PTA_High_ and PTA_Wide_) were also included as predictors to generate the Best-Subsets which take into account the effects of peripheral hearing loss. To avoid any spurious regression results caused by multicollinearity, subsets with variance inflation factors (VIFs) > 1.5 excluded (c.f., Stine, 1995). It is noteworthy that Best-Subset Regression is exploratory in nature and is often used when there is lack of a priori theory for a given topic. Here, it is not clear which neural parameter(s) can optimally model SPiN, and such an approach was used to answer this question and aid identification of the neural substrates that underlie SPiN.

After regression analyses using data from both young and older adults, the analyses were further conducted separately for the young and the older group, to evaluate whether they employ different neural mechanisms for SPiN.

## 3 Results

### 3.1 Behavioral results

ANOVA was conducted for SRT with Noise Type (SpN vs. BbN) and Age Group (young vs. older) as factors. The SRTs are plotted as boxplots^5^ in **Figure 4** for SpN and BbN of the 23 young and 18 older participants and the statistics are summarized in **Table 1**. The significant main effect of Noise Type (F_(1, 39)_ = 382.850, *p* < 10^−21^; SRT_SpN_ < SRT_BbN_) is consistent with previous finding that speech is better recognized in SpN than in BbN (Rosen et al., 2013). The significant main effect of Age Group (F_(1, 39)_ = 5.527, *p* = 0.024; SRT_Young_ < SRT_Older_) showed that young adults had better performance than older adults. A significant [Noise Type × Age Group] interaction occurred (F_(1, 39)_ = 10.010, *p* = 0.003) and post-hoc t-tests showed that young adults had significantly better performance than older adults in BbN (t_(26.667)_ = −3.399, *p* = 0.002, Cohen’s d = 1.132; equal variances not assumed), but not in SpN (t_(39)_ = −0.135, *p* = 0.893, Cohen’s d = 0.043; equal variances assumed).

**Table 1.** Statistical results of ANOVAs for SRT with factors of Noise Type (SpN vs. BbN) and Age Group (young vs. older). DV, df, F, and *p* refer to the dependent variable, degrees of freedom, F values, *p* values, respectively. Significant *p* values (< 0.05) are indicated in bold. * = *p* < 0.05; ** = *p* < 0.01; *** = *p* < 0.001.

| DV | Factors | *df1* | *df2* | *F* | *P* |
| --- | --- | --- | --- | --- | --- |
| SRT | Noise Type | 1 | 39 | 382.850 | **< 10<sup>-21</sup>\*\*\*** |
|  | Age Group | 1 | 39 | 5.527 | **0.024\*** |
|  | Noise Type × Age Group | 1 | 39 | 10.010 | **0.003\*\*** |

**Figure 4.**
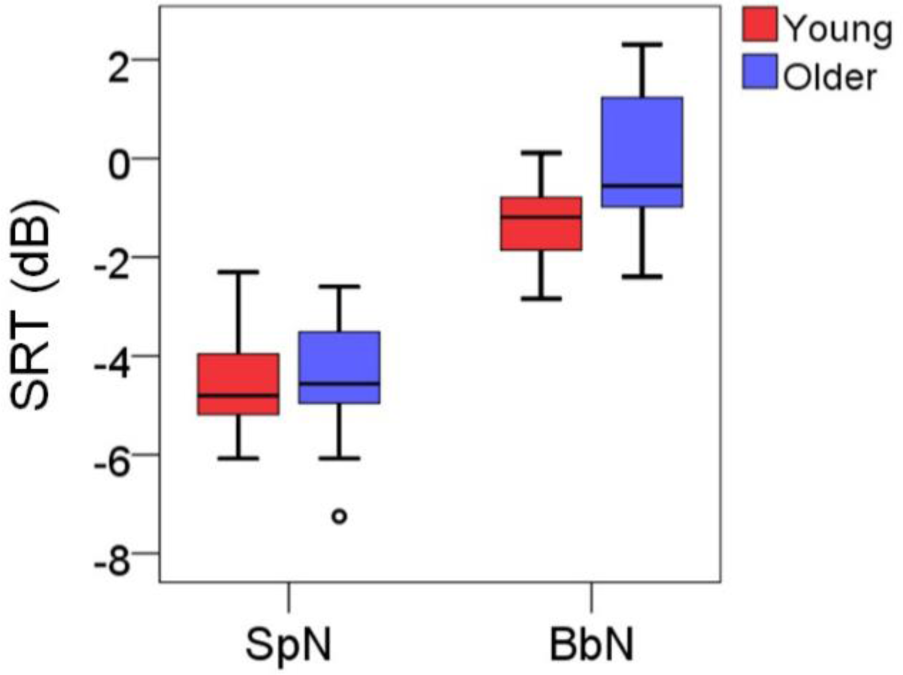
Boxplots for SRT as a function of Noise Type (SpN vs. BbN) and Age Group (young vs. older). Low SRTs represent good SPiN.

### 3.2 Neural results

Robustness of FFR was first confirmed by testing whether FFR magnitudes were statistically greater than their corresponding EEG noise floors (see *2.4.5*). It was shown that spectral magnitudes of FFRs were all significantly greater than noise floors (FFR_ENV_F0_Quiet_: *p* < 10^−8^; FFR_ENV_F0_SpN_ and FFR_ENV_F0_BbN_: *p* < 0.001; FFR_TFS_H2_Quiet_: *p* < 10^−11^; FFR_TFS_H2_SpN_ and FFR_TFS_H2_BbN_: *p* < 10^−^10; FFR_TFS_F2F3_Quiet_, FFR_TFS_F2F3_SpN_ and FFR_TFS_F2F3_BbN_: *p* < 10^−6^). Furthermore, the response signal-to-noise ratio (SNR, i.e., difference in magnitudes between FFRs and the corresponding noise floors) did not differ between age groups (SNR for FFR_ENV_F0_Quiet_: *p* > 0.07; SNRs for other FFR signatures: p > 0.2), indicating that SNR would not be a good index for measuring age differences. Additional simulations were also conducted in the present study showing that FFR magnitudes can more reliably quantify the FFR fidelity compared to response SNRs (see *Appendix 2*).

ANOVAs were then conducted for EEG signatures (FFR_ENV_F0_, FFR_PLV_F0_, FFR_TFS_H2_, FFR_TFS_F2F3_ and Logit-θ-PLV) including Noise Type (Quiet, SpN and BbN) and Age Group (young vs. older) as factors. Participants’ data were not included if artefact-free sweeps in the high arousal periods (wakefulness and nREM stage 1) were < 1450 in Quiet, SpN or BbN to ensure good EEG signal quality (see *2.4.4*). 29 (15) participants were retained (i.e., 12 (8) were rejected; rejection rate 29% where the numbers in brackets represent the numbers of young adults). Time series of the EEG signatures are given in **Figure 5** and the boxplots are shown in **Figure 6**.

**Figure 5.**
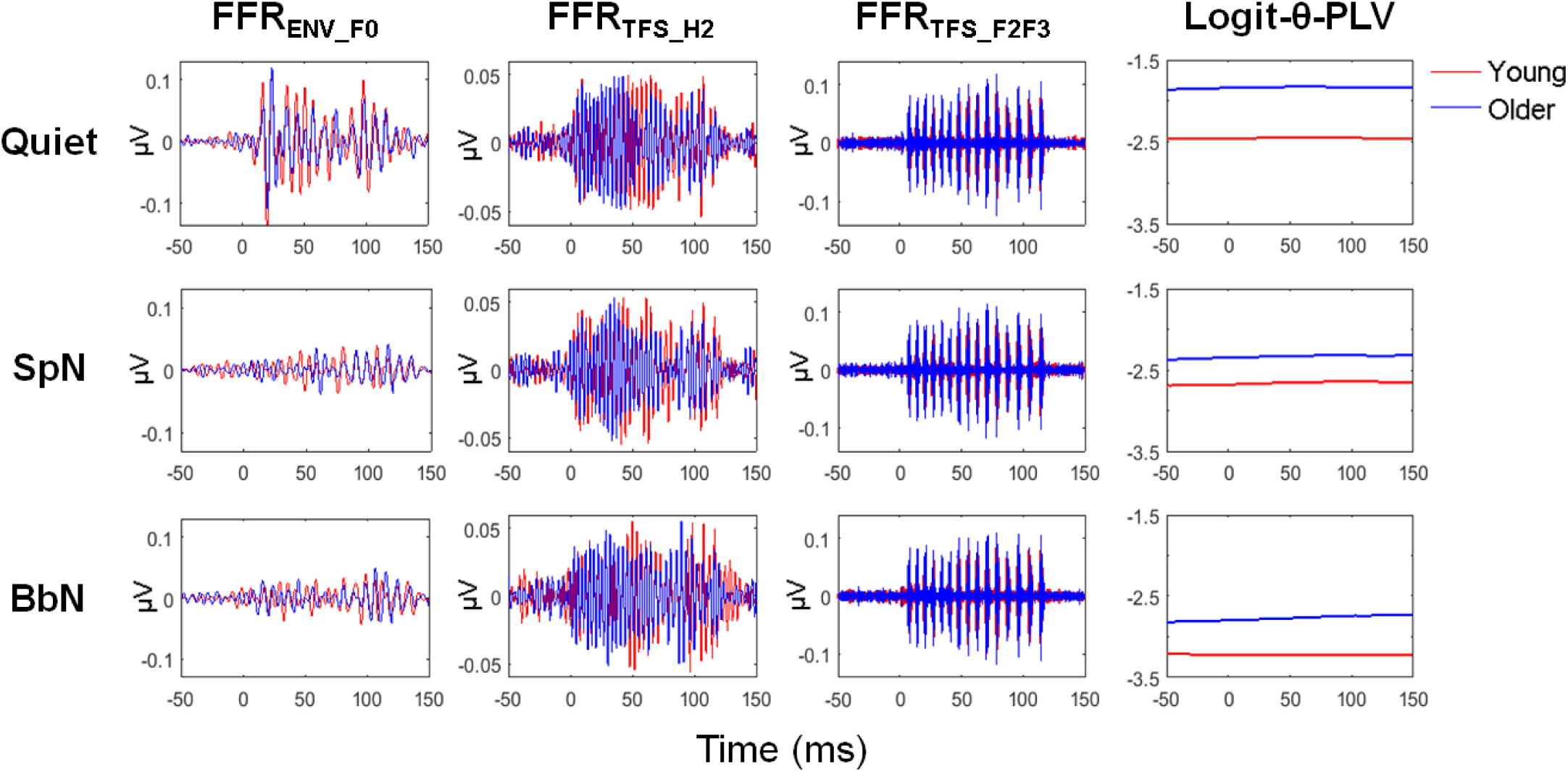
Time series of FFR_ENV_F0_, FFR_TFS_H2_, FFR_TFS_F2F3_ and Logit-θ-PLV in Quiet (upper row), SpN (mid row) and BbN (lower row). The series were based on sweeps with normalized numbers (1,450 ∼ 1,550) during the high arousal periods (see *2.4.3 and 2.4.4*) averaged across young (red) and older (blue) adults. Series of FFR_ENV_F0_ are shown as FFR_ENV_ bandpass filtered at 90 ∼ 180 Hz (corresponding to the F_0_ range). Series of FFR_TFS_H2_ and FFR_TFS_F2F3_ are shown as FFR_TFS_ bandpass filtered at 200 ∼ 340 Hz (corresponding to the H2 range), and at 2000 ∼ 4000 Hz (corresponding to the F2 and F3 range), respectively. ‘0’ represents the syllable onset.

**Figure 6.**
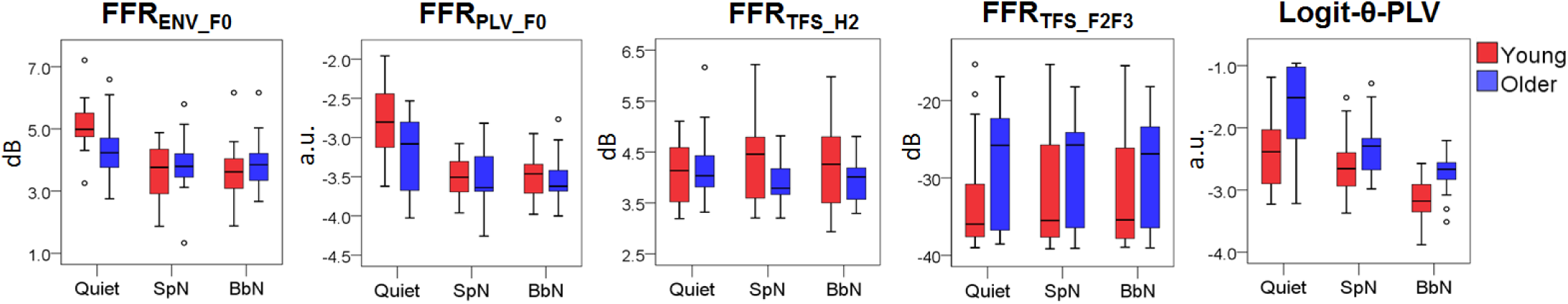
Boxplots for the five EEG signatures as a function of Noise Type (Quiet, SpN and BbN) and Age Group (young vs. older). FFR_ENV_F0_ and FFR_TFS_H2_ were measured as log-power (dB); FFR_TFS_F2F3_ was measured as power of cepstral spectrum (dB) at F2 and F3 range; FFR_PLV_F0_ and Logit-θ-PLV were measured as logit-transformed phase-locking values (a.u., arbitrary units). Young and older adults are plotted in red and blue respectively.

**Figure 7.**
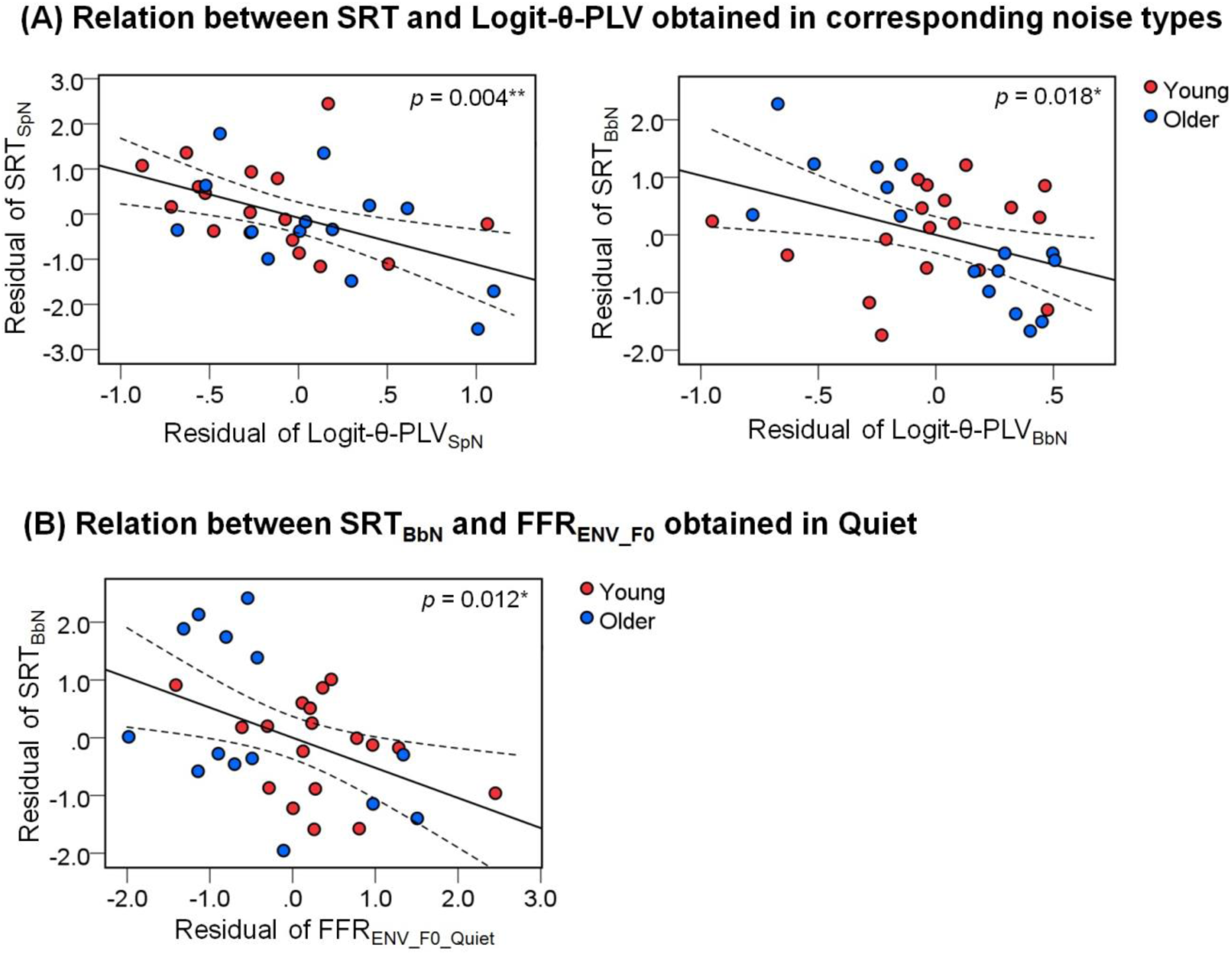
Scatter plots (young + older) for significant partial correlations after controlling for PTAs that visualize the relation between **(A)** SRT and Logit-θ-PLV obtained in the corresponding noise types (see statistics in **Table 4**); and **(B)** SRT_BbN_ and FFR_ENV_F0_ magnitude obtained in Quiet (see statistics in **Table 5**). Red and blue dots represent young and older participants, respectively.

Statistics on the ANOVAs are summarized in **Table 2**. Greenhouse-Geisser corrections were conducted to correct for violations of sphericity. For FFRs, significant main effects of Noise Type and [Noise Type × Age Group] interactions were found for both FFR_ENV_F0_ and FFR_PLV_F0_. Post-hoc comparisons following the main effects showed that FFR_ENV_F0_ and FFR_PLV_F0_ were significantly greater in Quiet than in noise (FFR_ENV_F0_Quiet_ > FFR_ENV_F0_SpN,_ *p* < 10^−7^, Cohen’s d = 1.297; FFRENV_F0_Quiet > FFRENV_F0_BbN, *p* < 10^−8^, Cohen’s d = 1.651; FFRPLV_F0_Quiet > FFRPLV_F0_SpN, *p* < 10^−6^, Cohen’s d = 1.201; FFR_PLV_F0_Quiet_ > FFR_PLV_F0_BbN,_ *p* < 10^−7^, Cohen’s d = 1.346), but they did not differ between SpN and BbN (both *p* > 0.5). Post-hoc t-tests following the interaction effects found significant greater FFR_PLV_F0_ for young than older adults in Quiet (t_(27)_ = −2.105, *p* = 0.045, Cohen’s d = 0.782). No main effects nor interactions were found for FFR_TFS_H2_ or FFR_TFS_F2F3_. For Logit-θ-PLV, there were significant main effects of Noise Type and Age Group, but no significant [Noise Type × Age Group] interaction. Post-hoc comparisons found that Logit-θ-PLV was greater in Quiet than in noise, and greater in SpN than in BbN (Logit-θ-PLV_Quiet_ > Logit-θ-PLV_SpN_, *p* = 0.002, Cohen’s d = 0.654; Logit-θ-PLV_Quiet_ > Logit-θ-PLV_BbN_, *p* < 10^−6^, Cohen’s d = 1,334; Logit-θ-PLV_SpN_ > Logit-θ-PLV_BbN_, *p* < 10^−4^, Cohen’s d = 0.961); Logit-θ-PLV was greater for older than young adults (*p* < 0.001, Cohen’s d = 1.504).

**Table 2.** Statistical results for ANOVAs (with Greenhouse-Geisser correction) for the five EEG signatures with factors of Noise Type (Quiet, SpN and BbN) and Age Group (young vs. older). DV, df, F, *p* refer to the dependent variable, degrees of freedom, F values, and *p* values (uncorrected), respectively. Significant *p* values (< 0.05) ae indicated in bold. ** = *p* < 0.01; *** = *p* < 0.001.

| DV | Factors | *df1* | *df2* | *F* | *p* |
| --- | --- | --- | --- | --- | --- |
| $FFR_{ENV\_F0}$ | Noise Type | 1.606 | 43.359 | 49.536 | **$< 10^{-10}$\*\*\*** |
|  | Age Group | 1 | 27 | 0.326 | 0.573 |
| | Noise Type × Age Group | 1.606 | 43.359 | 5.608 | **0.011\*** |
| $FFR_{PLV\_F0}$ | Noise Type | 1.559 | 42.082 | 42.827 | **$< 10^{-9}$\*\*\*** |
|  | Age Group | 1 | 27 | 2.225 | 0.147 |
| | Noise Type × Age Group | 1.559 | 42.082 | 3.522 | **0.049\*** |
| $FFR_{TFS\_H2}$ | Noise Type | 1.898 | 51.244 | 0.324 | 0.713 |
|  | Age Group | 1 | 27 | 0.005 | 0.946 |
| | Noise Type × Age Group | 1.898 | 51.244 | 2.061 | 0.140 |
| $FFR_{TFS\_F2F3}$ | Noise Type | 1.599 | 43.181 | 1.592 | 0.217 |
|  | Age Group | 1 | 27 | 2.479 | 0.127 |
| | Noise Type × Age Group | 1.599 | 43.181 | 0.051 | 0.918 |
| Logit-θ-PLV | Noise Type | 1.728 | 46.654 | 31.068 | **< 10<sup>-7</sup>\*\*\*** |
|  | Age Group | 1 | 27 | 16.386 | **< 0.001\*\*\*** |
| | Noise Type × Age Group | 1.728 | 46.654 | 0.972 | 0.375 |

ANCOVAs were conducted next that included PTAs (PTA_Low_, PTA_High_ and PTA_Wide_) as covariates. For FFR_ENV_F0_, an additional significant main effect of Age Group was found (F_(1, 26)_ = 8.842, *p* = 0.006, FFR_ENV_F0_Young_ > FFR_ENV_F0_Older_) compared to the previous ANOVA when PTA_Low_ was the covariate. The main effect of PTA_Low_ was also significant (F_(1, 26)_ = 12.943, *p* = 0.001), where higher PTA_Low_ correlated with greater FFR_ENV_F0_ magnitude (see **Table 3**). No significant main effects of Age or PTA were found when PTA_High_ or PTA_Wide_ were used as covariates (all *p* > 0.2) (see *Appendix* **Tables A2-A3**). For Logit-θ-PLV, the significant main effect of Age Group was maintained (F_(1, 26)_ = 4.608, *p* = 0.041 and F_(1, 26)_ = 5.937, *p* = 0.022) when PTA_Low_ and PTA_High_ were used as the covariates, respectively, however, this became non-significant (F_(1, 26)_ = 3.516, *p* = 0.072) when PTA_Wide_ was used as the covariate. No significant main effects of PTA_Low_, PTA_High_, or PTA_Wide_ occurred (all *p* > 0.2) (see *Appendix* **Tables A1-A3**). For FFR_PLV_F0_, FFR_TFS_H2_ and FFR_TFS_F2F3_, no significant main effects occurred (all *p* > 0.1) (see *Appendix* **Tables A1-A3**). The overall ANCOVA results showed: (1) FFR_ENV_F0_ declined with age when PTA_Low_ was controlled for, where higher PTA_Low_ was related to greater FFR_ENV_F0_; (2) the age effect for Logit-θ-PLV was maintained when PTAs were controlled for (though dropped below significance when PTA_Wide_ was used as the covariate) and the increased Logit-θ-PLV cannot be statistically explained by increased PTA.

**Table 3.** Statistical result of ANCOVA (with Greenhouse-Geisser correction) for FFR_ENV_F0_ with PTA_Low_ as the covariate. DV, df, F, and *p* refer to the dependent variable, degrees of freedom, F values, and *p* values (uncorrected), respectively. Significant *p* values (< 0.05) are indicated in bold. * = *p* < 0.05; ** = *p* < 0.01.

| DV | Factors | *df1* | *df2* | *F* | *p* |
| --- | --- | --- | --- | --- | --- |
| FFR<sub>ENV_F0</sub> | Noise Type | 1.506 | 39.152 | 50.277 | **< 10<sup>-9</sup>\*\*\*** |
|  | Age Group | 1 | 26 | 8.842 | **0.006\*\*** |
|  | PTA<sub>Low</sub> | 1 | 26 | 12.943 | **0.001\*\*** |
| | Noise Type × Age Group | 1.506 | 39.152 | 3.118 | 0.069 |
| | Noise Type × PTA<sub>Low</sub> | 1.506 | 39.152 | 1.445 | 0.246 |

### 3.3 Behavioral-neural relationship

Behavioral-neural relationships were assessed by linear regressions in which SRT in SpN was predicted by EEG signatures obtained in SpN, whilst SRT in BbN was predicted by EEG signatures obtained in BbN. SRTs were further predicted by EEG signatures obtained in Quiet. PTAs (PTA_Low_, PTA_High_ or PTA_Wide_) were also included as predictors provided that including them improved the statistical capacity of EEG signatures to predict SRTs in the Best-Subsets. Similar to the procedure in *3.2*, participants’ data obtained under a particular noise type (Quiet, SpN or BbN) were not included if artefact-free sweeps in the high arousal periods were < 1450 for that noise type (see *2.4.4*). This resulted in 31 (17) participants retained (i.e., 10 (6) were excluded; rejection rate 24%) for analyses in SpN, 34 (18) participants retained (i.e., 7 (5) were excluded; rejection rate 17%) for analyses in BbN, and 32 (18) participants retained (9 (5) were excluded; rejection rate 22%) for analyses in Quiet (the numbers in brackets represent the numbers of young adults).

#### 3.3.1 Regression results including data of both young and older adults

Statistics for the Best-Subset Regressions are shown as in **Tables 4 and 5**. Results that included data of both young and older adults were analyzed first. When SRTs were predicted by EEG signatures obtained in the respective noise types, SRTs were significantly correlated with Logit-θ-PLV (SpN, t_(28)_ = −3.104, *p* = 0.004; BbN, t_(31)_ = −2.508, *p* = 0.018; greater Logit-θ-PLV correlated with better SPiN) after PTA was controlled for (PTA_High_ for SpN and PTA_Wide_ for BbN) (**Table 4**). When SRTs were predicted by EEG signatures obtained in Quiet, a significant correlation was found between FFR_ENV_F0_ magnitude and SRT in BbN (t_(29)_ = −2.698, *p* = 0.012; greater FFR_ENV_F0_ magnitude correlated with better SPiN) after PTA_Low_ was controlled for (**Table 5**).

**Table 4.** Results for the Best-Subset Regressions in which SRTs were predicted by EEG signatures obtained in the corresponding noise types (i.e., SRT_SpN_ was predicted by EEGs obtained in SpN; SRT_BbN_ was predicted by EEGs obtained in BbN). DV refers to the dependent variables; *β*, CI, *T, p*, VIF refer to standardized *β*-coefficient, 95% confidence interval for standardized *β, t* values, *p* values (uncorrected) and variance inflation factors, respectively. *N* denotes the numbers of participants. *F* denotes the F values of the models (with corresponding *p* values in the brackets). Significant *p* values (< 0.05) are in bold. * = *p* < 0.05; ** = *p* < 0.01; *** = *p* < 0.001.

| DV | Participants | $N$ | Predictors | $\beta$ | CI | $T$ | $p$ | VIF | $F$ |
| --- | --- | --- | --- | --- | --- | --- | --- | --- | --- |
| SRT<sub>SpN</sub> | Young + older | 31 | Logit-θ-PLV<sub>SpN</sub> | -0.499 | [-0.828, -0.170] | -3.104 | **0.004\*\*** | 1.040 | 6.104 |
|  |  |  | PTA<sub>High</sub> | 0.351 | [0.022, 0.680] | 2.183 | **0.038\*** | 1.040 | **(0.006\*\*)** |
|  | Young | 17 | PTA<sub>High</sub> | 0.445 | [-0.048, 0.938] | 1.922 | 0.074 | 1.000 | 3.695 (0.074) |
|  | Older | 14 | FFR<sub>ENV_F0_SpN</sub> | -0.475 | [-0.903, -0.048] | -2.480 | **0.033\*** | 1.042 | 6.120 |
| | | | Logit-θ-PLV<sub>SpN</sub> | -0.475 | [-0.903, -0.047] | -2.475 | **0.033\*** | 1.045 | **(0.012\*)** |
|  |  |  | PTA<sub>Wide</sub> | 0.363 | [-0.064, 0.791] | 1.892 | 0.088 | 1.045 |  |
| SRT<sub>BbN</sub> | Young + older | 34 | Logit-θ-PLV<sub>BbN</sub> | -0.377 | [-0.684, -0.070] | -2.508 | **0.018\*** | 1.365 | 14.738 |
|  |  |  | PTA<sub>Wide</sub> | 0.814 | [0.506, 1.122] | 5.418 | **< 10<sup>-5</sup>\*\*\*** | 1.365 | **(< 10<sup>-4</sup>\*\*\*)** |
|  | Young | 18 | PTA<sub>Wide</sub> | 0.318 | [-0.184, 0.820] | 1.343 | 0.198 | 1.000 | 1.804 (0.198) |
|  | Older | 16 | FFR<sub>TFS_H2_BbN</sub> | -0.351 | [-0.578, -0.124] | -3.368 | **0.006\*\*** | 1.047 | 28.054 |
| | | | Logit-θ-PLV<sub>BbN</sub> | -0.769 | [-1.005, -0.533] | -7.103 | **< 10<sup>-4</sup>\*\*\*** | 1.128 | **(< 10<sup>-4</sup>\*\*\*)** |
|  |  |  | PTA<sub>Wide</sub> | 0.696 | [0.456, 0.936] | 6.350 | **< 10<sup>-4</sup>\*\*\*** | 1.157 |  |

**Table 5.** Results for the Best-Subset Regression in which SRTs (both in SpN and BbN) were predicted by EEG signatures obtained in Quiet. DV refers to the dependent variables; *β*, CI, *T, p*, VIF refers to standardized *β*-coefficient, 95% confidence interval for standardized *β, t* values, *p* values (uncorrected) and variance inflation factors, respectively. *N* denotes the numbers of participants. *F* denotes the F values of the models (with corresponding *p* values in the brackets). Significant *p* values (< 0.05) are in bold. * = *p* < 0.05; ** = *p* < 0.01.

| DV | Participants | $N$ | Predictors | $\beta$ | CI | $T$ | $p$ | VIF | $F$ |
| --- | --- | --- | --- | --- | --- | --- | --- | --- | --- |
| SRT<sub>SpN</sub> | Young + older | 32 | FFR<sub>ENV_F0_Quiet</sub> | -0.307 | [-0.661, 0.048] | -1.765 | 0.088 | 1.000 | 3.116 (0.088) |
|  | Young | 18 | FFR<sub>ENV_F0_Quiet</sub> | -0.508 | [-1.001, -0.014] | -2.192 | **0.045\*** | 1.161 | 3.304 (0.065) |
|  |  |  | PTA<sub>Wide</sub> | 0.478 | [-0.016, 0.973] | 2.062 | 0.057 | 1.161 |  |
|  | Older | 14 | FFR<sub>ENV_F0_Quiet</sub> | -0.471 | [-1.010, 0.069] | -1.919 | 0.081 | 1.068 | 3.375 (0.072) |
|  |  |  | FFR<sub>TFS_H2_Quiet</sub> | -0.534 | [-1.074, 0.005] | -2.179 | 0.052 | 1.068 |  |
| SRT<sub>BbN</sub> | Young + older | 32 | FFR<sub>ENV_F0_Quiet</sub> | -0.410 | [-0.720, -0.099] | -2.698 | **0.012\*** | 1.012 | 7.425 |
|  |  |  | PTA<sub>Low</sub> | 0.460 | [0.150, 0.771] | 3.030 | **0.005\*\*** | 1.012 | **(0.002\*\*)** |
|  | Young | 18 | PTA<sub>Wide</sub> | 0.323 | [-0.181, 0.829] | 1.363 | 0.192 | 1.000 | 1.858 (0.192) |
|  | Older | 14 | FFR<sub>ENV_F0_Quiet</sub> | -0.455 | [-0.857, -0.054] | -2.525 | **0.030\*** | 1.069 | 7.622 |
|  |  |  | FFR<sub>TFS_H2_Quiet</sub> | -0.551 | [-0.969, -0.133] | -2.941 | **0.015\*** | 1.153 | **(0.006\*\*)** |
|  |  |  | PTA<sub>Wide</sub> | 0.441 | [0.037, 0.845] | 2.434 | **0.035\*** | 1.081 |  |

#### 3.3.2 Regression results in the young and the older group separately

Best-Subset Regressions were then conducted separately for the young and older groups. In the young group, SRT in SpN was significantly correlated with FFR_ENV_F0_ magnitude obtained in Quiet after PTA_Low_ was controlled for (t_(15)_ = −2.195, *p* = 0.045; greater FFR_ENV_F0_ magnitude correlated with better SPiN; **Table 5**). No significant correlations were found between SRTs and EEG signatures in the corresponding noise types nor between SRT in BbN and in EEG signatures obtained in Quiet (**Tables 4** and **5**). In the older group, when SRTs were predicted by EEG signatures in the respective noise types, SRTs were significantly correlated with FFRs (FFR_ENV_F0_ for SpN: t_(10)_ = −2.480, *p* = 0.033; FFR_TFS_H2_ for BbN: t_(12)_ = −3.368, *p* = 0.006) and Logit-θ-PLV (SpN: t_(10)_ = −2.475, *p* = 0.033; BbN: t_(12)_ = −7.103, *p* < 10^−4^) (greater FFR magnitudes and Logit-θ-PLV correlated with better SPiN) after PTA_Wide_ was controlled for (**Table 4**). When SRTs were predicted by EEG signatures in Quiet, SRT in BbN was significantly correlated with FFR_ENV_F0_ and FFR_TFS_H2_ (FFR_ENV_F0_: t_(10)_ = −2.941, *p* = 0.030; FFR_TFS_H2_: t_(10)_ = −2.525, *p* = 0.015; greater FFR magnitudes correlated with better SPiN) after PTA_Wide_ was controlled for (**Table 5**).

Taken together, the regression analyses showed that: (1) when combining data from both young and older adults, by controlling for the degree of hearing loss (PTAs), SRTs can be predicted by cortical phase-locked responses (Logit-θ-PLV) to speech obtained in noise (greater Logit-θ-PLV associated with better SPiN) and by subcortical phase-locked responses to speech F_0_ obtained in Quiet (greater FFR_ENV_F0_ magnitude associated with better SPiN); (2) SRTs are predicted by subcortical and cortical responses (FFR_ENV_F0_, FFR_TFS_H2_ and Logit-θ-PLV) obtained in noise in the older group, not in the young group, and by subcortical responses obtained in Quiet (FFR_ENV_F0_/FFR_TFS_H2_) in the young (SRT_SpN_) and the older group (SRT_BbN_).

### 3.4 Cortical-subcortical/peripheral relationship

Correlations between Logit-θ-PLV and FFR signatures were conducted across the young and older adults to assess the cortical-subcortical/peripheral relationship in different noise types (Quiet, SpN and BbN). This was to see how such relationships are linked to SPiN. It was found that Logit-θ-PLV showed significant positive correlations with FFR_TFS_ magnitudes in SpN (N = 31; with FFR_TFS_H2_, r = 0.441; *p* = 0.013; and with FFR_TFS_F2F3_, r = 0.560, *p* = 0.001). As Logit-θ-PLV was significantly associated with SRT in SpN (see **Table 4**), FFR_TFS_ may then indirectly impact SPiN. No significant correlations were found between Logit-θ-PLV and FFR_TFS_ in other noise types (Quiet or BbN), nor between Logit-θ-PLV and FFR_ENV_F0_/FFR_PLV_F0_ in any noise type, even after age or PTAs were controlled for.

## 4 Discussion

The relationships between age-related phase-locked responses to speech in the auditory sensory systems and SPiN were investigated. Participant groups were young and older adults. The older adult group had peripheral hearing at high frequencies that ranged from normal to mild/moderate hearing loss. Such demographics are typical in normal aging populations (Popinath et al., 2009; Humes et al., 2010). Other studies have looked specifically at normal-hearing older adults (Anderson et al., 2011, 2012; Presacco et al., 2016a; Schoof and Rosen, 2016).

Different patterns of age effects on auditory phase-locked responses were revealed at the subcortical and cortical levels: aging is related to decreases in subcortical responses (FFRs) and increases in cortical responses (θ-PLV). Relationships between behavioral and neural performance showed that subcortical responses obtained in quiet and cortical responses obtained in noise had significant positive associations with SPiN, indicating that effects of aging on subcortical (decrease with aging) and cortical (increase with aging) activities make different impacts on SPiN.

### 4.1 Age effect on subcortical phase-locked activity (FFRs)

Speech-evoked FFRs originate primarily from the brainstem (Chandrasekaran and Kraus, 2010; Bidelman, 2018). Previous studies, including ones on older adults who had normal audiometric hearing, reported that FFRs declined with age (Anderson et al., 2012; Presacco et al., 2016a). On the other hand, evidence shows that hearing loss can reduce neural inhibition that allows greater encoding of F_0_-rate envelope modulations in both animals (Kale and Heinz, 2010; Henry et al., 2014; Zhong et al., 2014) and humans (Anderson et al., 2013; Goossens et al., 2019).

In the present study, no significant differences were found between young and older adults for magnitudes of FFRs (FFR_ENV_F0_, FFR_TFS_H2_, or FFR_TFS_F2F3_). However, additional ANCOVAs showed that FFR_ENV_F0_ magnitude was significantly smaller in older than young adults after PTA_Low_ (0.25 ∼ 1 kHz) was controlled for. Also, PTA_Low_ correlated positively with the FFR_ENV_F0_ magnitude (i.e., FFR_ENV_F0_ magnitude increased with low-frequency hearing loss). Therefore, these results are in line with findings that encoding of envelopes at high-gamma frequencies corresponding to the F_0_ range declines during aging when peripheral hearing is normal but increases when there is hearing loss (Goossens et al., 2016, 2019), indicating the age-related neural declines in phase-locking and reduced neural inhibition related to hearing loss.

However, caution is required when interpreting such results. FFR magnitudes are related to participants’ sensitivity to the syllable where greater audibility leads to higher magnitudes (Ananthakrishnan et al., 2016). Therefore, the relation between PTA and FFR magnitudes probably depends on the individual’s degree of hearing loss. This could possibly explain why PTA_Low_, but not PTA_High_, correlated with FFR magnitude, as the older adults in the present study had a much smaller range of PTA_Low_ (< 30 dB HL) than PTA_High_ (from normal hearing to mild/moderate hearing loss) (see **Figure 1**) and the wider range of PTA_High_ may lead to a greater mix of reduced audibility and neural inhibition on FFRs. Taken together, effects of aging on subcortical phase-locked processing may be due to a combined effect of age-related hearing loss and the aging process that is independent of peripheral hearing.

### 4.2 Age effect on cortical phase-locked activity (θ-PLV)

θ-PLV was calculated as low-frequency (theta-band) phase-locked activity time-locked to the syllable onset. It reflects the tracking of speech envelopes (Luo and Poeppel, 2007; Howard and Poeppel, 2010; Peelle et al., 2013) and neural excitability in response to acoustic stimuli in the auditory cortex (Ng et al., 2013). The present study found that θ-PLV increased with age, consistent with previous studies (Tlumak et al., 2015; Goossens et al., 2016). This is also in line with findings that older adults have larger magnitudes of cortical auditory-evoked responses (Alain et al., 2014; Herrmann et al., 2013, 2016) and greater cortical tracking of speech envelopes (Presacco et al., 2016a).

Further ANCOVAs showed that, after PTAs (PTA_Low_ and PTA_High_) were controlled for, the statistical effect of age was maintained. No statistical correlations occurred between PTAs and θ-PLV in ANCOVAs, indicating that the age-related increase cannot be explained by hearing loss. Consequently, the results may be attributable to hyperexcitiblity of the central auditory system during the aging process (Caspary et al., 2008). Note that in the present study, participants listened passively to a repeatedly-presented syllable that lacked higher-order semantic information. This indicates that the age effect may occur at an early stage in the auditory cortex.

It is noteworthy that the effects of age and PTAs on FFRs and θ-PLV differed, i.e., FFR_ENV_F0_ declined with age when PTA_Low_ was controlled for, whereas θ-PLV increased with age; FFR_ENV_F0_ increased with PTA_Low_, whereas no association occurred between θ-PLV and PTAs. Thus these results imply that the mechanisms by which aging and peripheral hearing loss affect the phase-locked neural activities differ at subcortical and cortical levels.

### 4.3 Contributions of age effects on subcortical and cortical activities to SPiN

The central question for the present study concerned how the effects of age on phase-locked activities contributed to impaired SPiN. This question can be approached by conducting regression analyses on participants’ data across a wide range of ages. Here, linear regressions were conducted to study the neural-behavioral relationship using combined data from young and older adults, as well as young and older adults separately. The present study used an approach that differs from previous attempts (Presacco et al., 2016a; Goossens et al., 2018). Presacco et al. (2016a) studied the relation between speech-evoked phase-locked responses (FFRs and cortical tracking of speech envelopes) and SPiN in both young and older adults, but did not find any significant neural-SPiN correlations. A limitation in this study was that different types of background noise were used in the neural recording (single-talker background) and SPiN tasks (four-talker babble). Hence the neural-SPiN association was not appropriately assessed because acoustic and linguistic properties differed between the two types of background noise (see the discussion in Presacco et al., 2016). Goossens et al., (2018) investigated the relation between subcortical/cortical auditory steady-state responses (ASSRs) and SPiN in both normal-hearing and hearing impaired adults across ages and included age itself as an additional predictor when modeling the neural-behavioral relation. This did not fulfil the aim of testing how age-related neural factors contribute to SPiN. In the present study, regressions were conducted with neural signatures and SPiN under the same types of background noise. That is, SPiN in SpN and 16-talker BbN were predicted by neural data obtained in SpN and 16-talker BbN, respectively; neural data obtained in quiet were additionally used as predictors, as FFRs in quiet could be associated with SPiN (Anderson et al., 2011). Also, age itself was not included as a predictor.

Previous studies have argued that declines in subcortical speech-evoked FFRs are an important determinant of SPiN difficulty in older adults (Anderson et al., 2011, 2012; Presacco et al., 2016a). Consistent with this argument, the present study showed the FFR_ENV_F0_ declines with age and that decreased FFR _ENV_F0_ obtained in quiet was associated with decreased SPiN. At the cortical level, θ-PLV, which was shown to increase with age, was associated with increased SPiN. Thus this argues against the view that increased tracking of speech envelopes or auditory cortical excitability reflects a diminished excitation-inhibition balance in the neural network which results in impaired SPiN (Presacco et al., 2016a). An alternative explanation may be that, cortical hyperexcitability *per se* does not impair SPiN, but influences SPiN in an indirect way: increased cortical excitability is significantly related to decreased attention and inhibitory control in older adults which could, in turn, impair SPiN (Presacco et al., 2016b). The current results thus showed that impacts of age effects on SPiN are different at subcortical (decreased FFR_ENV_F0_ with age associated with decreased SPiN) and cortical (increased θ-PLV with age associated with increased SPiN) levels.

The results also showed potential different mechanisms of neural-behavioral relationships across ages, in which SPiN is better predicted by cortical and subcortical signatures obtained in noise in the older compared to the young group. This could be because higher-level cognitive functions, such as working memory and attention ability (not tested in the present study) that decline with age (see *4.4.2* for further discussions). These factors could play greater roles in modelling the individual variability of SPiN ability in young adults. The results could also arise because individual variability of SPiN was relatively small in young adults (especially SRT_BbN_, which had significantly lower variability in the young than the older group according to Levene’s test, see *3.1*). This would make SPiN more difficult to predict using linear regression.

Another interesting finding was that cortical, compared to subcortical, responses obtained in noise is a more reliable predictor of SPiN, which is reflected not only in regression analyses, but also by comparisons across noise types. It was shown that SPiN was significantly better in SpN than in BbN, consistent with previous work (Rosen et al., 2013). Correspondingly, θ-PLV was also significantly higher in SpN than in BbN. The explanation may be that, compared to SpN, BbN is an informational masker that causes phonetic interference (Rosen et al., 2013) leading to worse cortical envelope tracking of target speech, hence worse SPiN in BbN than in SpN. On the other hand, no such effect was found for FFRs when SpN was compared with BbN. It could be that, cortical, but not subcortical, phase-locked responses reflect the mechanisms underlying speech perception as a function of noise types. Alternatively, signal quality of FFRs (FFR_ENV_F0_) dropped significantly from quiet to SpN/BbN (see **Figures 5** and **6**), which could hinder its predictability for SPiN in different noise types. This might also explain why FFR_ENV_F0_ obtained in quiet, compared to that obtained in noise, showed better predictability (although FFR_ENV_F0_SpN_ also significantly predicted SRT_SpN_ in the older group, see **Table 4**).

### 4.4 Limitations and future work

#### 4.4.1 Participant samples

The present study recruited older participants who covered a wide range of peripheral hearing from normal to mild/moderate hearing loss. Although the degree of hearing loss was controlled for by using PTAs as covariates during the analyses, a more direct and better approach would be to compare normal-hearing older adults with those who had hearing loss to clarify the respective effects of aging and hearing loss. However, sample size in the present study (< 17 older participants after data rejection) made it difficult to use this approach. Future research will need to recruit participants with bigger sample sizes with better control in hearing loss for older adults.

#### 4.4.2 Possible role of higher-level functions for SPiN

Phase-locked activities were obtained in the present study with a paradigm where participants listened passively to repeated syllables without high-level linguistic features (semantic or syntactic information), hence reflecting early-stage sensory processing of speech in the auditory brainstem and cortex. Therefore, the present study focused mainly on how effects of age on sensory processing influenced SPiN, while higher-level cognitive functions were not measured.

The present results showed that older adults had significantly worse SPiN than young adults in BbN, but not in SpN, consistent with previous findings (Helfer and Freyman, 2008; Schoof and Rosen, 2014). However, no significant [Noise Type × Age] interactions were found for any of the EEG signatures, hence no sensory neural evidence was provided to explain this behavioral phenomenon. This leaves the alternative explanation through age-related declines in higher-level cognitive functions. Aging and age-related hearing loss are associated with declines in cognitive functions, such as working memory (WM) and attention (Lin et al., 2013). WM capacity is related to SPiN in older adults (Schoof and Rosen, 2014), while attention is critical for suppressing neural processing of background noise in SPiN (Rimmele et al., 2015) and such ability deteriorates during aging (Andres et al., 2006; Presacco et al., 2019). WM and attention may be particularly important for SPiN in BbN as they contribute to resisting the informational masking caused by BbN (Schneider et al., 2007; Shinn-Cunningham and Best, 2008).

As well as WM and attention capacity during SPiN, Mild Cognitive Impairment (MCI) could occur in some participants during normal aging (Petersen et al., 1999) and speech-evoked cortical and subcortical activities have been reported to be related to MCI that may have further affected SPiN (Anderson et al., 2013; Bidelman et al., 2017). For example, Bidelman et al. (2017) found that people with MCI showed hyperexcitability of subcortical and cortical responses to speech compared to controls. It is thus not clear how MCI contributes to the neural-behavioral relation found the in the present study. Future studies should include measurements of cognitive functions and screening for cognitive impairment in the model to better predict SPiN.

In addition, previous studies have shown that older adults have diminished sensitivity to temporal fine structure (TFS) that leads to degraded SPiN (Hopkins and Moore, 2011; Fullgrabe et al., 2015). However, although TFS processing at the resolved harmonics region (FFR_TFS_H2_) was significantly associated with SPiN in older adults (see **Table 4**, modelling SRT_BbN_), the present study did not find statistical evidence for age effects on TFS processing (FFR_TFS_H2_ or FFR_TFS_F2F3_). Diminished sensitivity to TFS in SPiN in older adults may at least in part arise from declines in TFS processing that involves higher-level cognitive function, compared to declines at early-stage sensory processing. Such claims may be further tested in the future.

#### 4.4.3 Paradigms used for EEG recording

In the present study, EEGs were recorded when participants listened to a repeatedly presented vowel over a loudspeaker. The reason for using a loudspeaker rather than inserted earphones was because the present older adult’s data were obtained from our previous study (Mai et al., 2018). This study included hearing-aid users as participants, for whom inserted earphones are not appropriate (their data, however, were not included here, see *Footnote 1* for detailed explanations). Changes in head and body positions were controlled by instructions to avoid movements and monitored by a webcam. However, the possibility of slight movements, such as head jitters, cannot be ruled out. Such possibilities can change the ear position and thus interfere with phase-locked responses to high-frequency TFS properties (FFR_TFS_H2_ and FFR_TFS_F2F3_) in the free field. Future research using inserted earphones thus is needed to confirm the current results relevant to FFR_TFS_.

Furthermore, although arousal effects were controlled in the present study by restricting the data used in analyses to those when participants were in the high arousal states, amount of attention to the acoustic stimuli was not controlled during the EEG recording. While selective auditory attention can modulate auditory cortical (Choi et al., 2013; Kong et al., 2014) and subcortical (Galbraith et al., 2003; Hairston et al., 2013; Lehmann and Schönwiesner, 2014) electrophysiological responses, it is not totally clear how they are affected by unpredictable changes in attention during passive listening as in the present study. Therefore, additional tasks of active listening to target speech stimuli under the corresponding noise types need to be conducted in the future to investigate whether the current neural-behavioral relationships is replicable.

#### 4.4.4 Cortical-subcortical interaction

Relationships between cortical and subcortical/peripheral responses were also tested. It was found that Logit-θ-PLV had significant positive correlations with FFR_TFS_H2_ and FFR_TFS_F2F3_ magnitudes in SpN, indicating that FFR_TFS_ might indirectly contribute to SPiN via cortical-subcortical interactions (see *3.4*). However, as they are correlations based on data across different participants, it is not known whether such correlations reflect cortical-subcortical connectivity within individuals, whether connectivity is afferent or efferent, and how the connectivity is associated with age or SPiN. It is necessary in future work to develop effective methods to quantify the strength and direction of cortical-subcortical interactions (directional phase relations between cortical responses and FFRs, e.g., Bidelman et al., 2019).

#### 4.4.5 Other factors that may influence the hearing loss

Degree of hearing loss was tested via air-conduction in the present study. The audiogram for majority of the participants showed hearing loss at high frequencies (≥ 2 kHz) (**Figure 1**), consistent with the pattern of presbycusis which is usually sensorineural hearing disorders caused by age-related functional declines in the inner ear. Despite this, we cannot totally exclude that some of the participants may also suffer from conductive hearing loss. Future work should further include bone-conduction to test for conductive hearing loss in order to better clarify the roles of different types of hearing loss.

### 4.5 Conclusions

Subcortical (FFRs) and cortical (θ-PLV) phase-locked activities reflect how the brain processes fine-grained acoustic properties (F_0_, temporal fine structures and acoustic envelopes) that are crucial for SPiN. The present study hypothesized that effects of age on these activities should be associated with SPiN. Compared to young adults, it was found that older adults have smaller FFR magnitude (when low-frequency hearing loss was controlled for) and greater θ-PLV, illustrating distinct mechanisms of age effects at the subcortical and cortical levels. Smaller FFR magnitude reflects declines in subcortical phase-locking during aging and was associated with decreased SPiN, whilst greater θ-PLV reflects the neural hyperexcitability in the auditory cortex during aging and was associated with increased SPiN. The current study thus provided evidence for different mechanisms at the sensory subcortical and cortical levels by which age affects speech-evoked phase-locked activities and SPiN. Future work need to be conducted by combining cognitive assessments to study how higher-level cognitive functions influence such mechanisms and contribute to SPiN together with sensory processing during aging.

## Supporting information

Appendix 1 ANCOVA results and Appendix 2 Simulations that test relative reliability of FFR measurements

## Conflict of Interest

The authors declare no competing conflicts of interest.

## Footnotes

1 The present study used the data of non-hearing-aid older adults in Group 2 of Mai et al. (2018), where participants listened to the same acoustic stimuli as in the present study whilst neural recordings were made. Data from hearing aid users in Mai et al. (2018) were not included here, since PTAs could not be measured precisely in these participants and hearing aids may introduce additional effects.

2 Since the spectral distribution in the speech stimulus used in both behavioral and neural assessments extended to ≤ 4 kHz, PTAs at 6 and 8 kHz were not included in the statistical analyses in the present study.

3 Three older participants had PTAs that were higher than the measurable limit of the audiometer (85 dB) at 8 kHz in either the left or the right ear and thresholds for them were set at 85 dB when calculating averages over both ears.

4 Lower rejection thresholds were used than with FFRs (± 25 μV) because the θ-band signal usually does not have excessively high amplitude since it occupies a relatively narrow frequency range. More than 80% of the sweeps were retained in all participants after artefact rejection.

5 N.B., circles in boxplots represent outliers which are defined as data points that fall outside the 1.5 times the interquartile range i.e. are above/below the upper/lower quartile. Outliers were *not* excluded during the statistical analyses.

## References

Aiken SJ, Picton TW. Envelope and spectral frequency-following responses to vowel sounds. Hearing Research. 2008; 245:35–47.

Alain C. Effects of age-related hearing loss and background noise on neuromagnetic activity from auditory cortex. Frontiers in Systems Neuroscience. 2014; 8:8.

Ananthakrishnan S, Krishnan A, Bartlett E. Human frequency following response: neural representation of envelope and temporal fine structure in listeners with normal hearing and sensorineural hearing loss. Ear and Hearing. 2016; 37:e91.

Anderson S, Parbery-Clark A, Yi HG, Kraus N. A neural basis of speech-in-noise perception in older adults. Ear and Hearing. 2011; 32:750.

Anderson S, Parbery-Clark A, White-Schwdoch T, Kraus N. Aging affects neural precision of speech encoding. Journal of Neuroscience. 2012; 32:14156–64.

Anderson S, White-Schwoch T, Parbery-Clark A, Kraus N. A dynamic auditory-cognitive system supports speech-in-noise perception in older adults. Hearing Research. 2013; 300:18–32.

Andrés P, Parmentier FB, Escera C. The effect of age on involuntary capture of attention by irrelevant sounds: a test of the frontal hypothesis of aging. Neuropsychologia. 2006; 44:2564–8.

Aviyente S, Bernat EM, Evans WS, Sponheim SR. A phase synchrony measure for quantifying dynamic functional integration in the brain. Human Brain Mapping. 2011; 32:80–93.

Bench J, Kowal Å, Bamford J. The BKB (Bamford-Kowal-Bench) sentence lists for partially-hearing children. British journal of audiology. 1979; 13:108–12.

Bidelman GM. Subcortical sources dominate the neuroelectric auditory frequency-following response to speech. NeuroImage. 2018; 175:56–69.

Bidelman GM, Price CN, Shen D, Arnott SR, Alain C. Afferent-efferent connectivity between auditory brainstem and cortex accounts for poorer speech-in-noise comprehension in older adults. Hearing research. 2019; 107795:1–12.

Bidelman GM, Lowther JE, Tak SH, Alain C. Mild cognitive impairment is characterized by deficient brainstem and cortical representations of speech. Journal of Neuroscience. 2017; 37(13):3610–20.

Boersma P, Weenink D. Praat: doing phonetics by computer [Computer program] Version 5.3.51, Retrieved 2 June 2013.

Burnham KP, Anderson DR. Model selection and multimodel inference: a practical information-theoretic approach. 2003; Springer Science & Business Media; 2003.

Carpenter AL, Shahin AJ. Development of the N1–P2 auditory evoked response to amplitude rise time and rate of formant transition of speech sounds. Neuroscience Letters. 2013; 544:56–61.

Caspary DM, Ling L, Turner JG, Hughes LF. Inhibitory neurotransmission, plasticity and aging in the mammalian central auditory system. Journal of Experimental Biology. 2008; 211:1781–91.

Chandrasekaran B, Kraus N. The scalp-recorded brainstem response to speech: Neural origins and plasticity. Psychophysiology. 2010; 47:236–46.

Choi I, Rajaram S, Varghese LA, Shinn-Cunningham BG. Quantifying attentional modulation of auditory-evoked cortical responses from single-trial electroencephalography. Front Hum Neurosci. 2013; 7:115

Czisch M, Wehrle R, Kaufmann C, Wetter TC, Holsboer F, Pollmächer T, Auer DP. Functional MRI during sleep: BOLD signal decreases and their electrophysiological correlates. European Journal of Neuroscience. 2004; 20:566–74.

Dajani HR, Purcell D, Wong W, Kunov H, Picton TW. Recording human evoked potentials that follow the pitch contour of a natural vowel. IEEE Transactions on Biomedical Engineering. 2005; 52:1614–8.

Fujihira H, Shiraishi K. Correlations between word intelligibility under reverberation and speech auditory brainstem responses in elderly listeners. Clinical Neurophysiology. 2015; 126:96–102.

Füllgrabe C, Moore BC, Stone, MA. Age-group differences in speech identification despite matched audiometrically normal hearing: contributions from auditory temporal processing and cognition. Frontiers in Aging Neuroscience. 2010; 6:347.

Galbraith GC, Olfman DM, Huffman TM. Selective attention affects human brain stem frequency-following response. Neuroreport. 2003;14:735–8.

Goossens T, Vercammen C, Wouters J, Wieringen AV. Aging affects neural synchronization to speech-related acoustic modulations. Frontiers in Aging Neuroscience. 2016; 8:133.

Goossens T, Vercammen C, Wouters J, van Wieringen A. Neural envelope encoding predicts speech perception performance for normal-hearing and hearing-impaired adults. Hearing Research. 2018; 370:189–200.

Goossens T, Vercammen C, Wouters J, van Wieringen A. The association between hearing impairment and neural envelope encoding at different ages. Neurobiology of Aging. 2019; 74:202–12.

Gopinath B, Rochtchina E, Wang JJ, Schneider J, Leeder SR, Mitchell P. Prevalence of age-related hearing loss in older adults: Blue Mountains Study. Archives of internal medicine. 2009; 169:415–8.

Hairston WD, Letowski TR, McDowell K. Task-related suppression of the brainstem frequency following response. PLoS One. 2013; 8:e55215.

Harris KC, Eckert MA, Ahlstrom JB, Dubno JR. Age-related differences in gap detection: Effects of task difficulty and cognitive ability. Hearing research. 2010; 264:21–9.

Helfer KS, Freyman RL. Aging and speech-on-speech masking. Ear and Hearing. 2008; 29:87.

Henry KS, Kale S, Heinz MG. Noise-induced hearing loss increases the temporal precision of complex envelope coding by auditory-nerve fibers. Frontiers in Systems Neuroscience. 2014; 8:20.

Herrmann B, Henry MJ, Scharinger M, Obleser J. Auditory filter width affects response magnitude but not frequency specificity in auditory cortex. Hearing Research. 2013; 304:128–36.

Herrmann B, Henry MJ, Johnsrude IS, Obleser J. Altered temporal dynamics of neural adaptation in the aging human auditory cortex. Neurobiology of Aging. 2016; 45:10–22.

Hopkins K, Moore BC. The effects of age and cochlear hearing loss on temporal fine structure sensitivity, frequency selectivity, and speech reception in noise. The Journal of the Acoustical Society of America. 2011; 130:334–49.

Howard MF, Poeppel D. Discrimination of speech stimuli based on neuronal response phase patterns depends on acoustics but not comprehension. Journal of neurophysiology. 2010; 104:2500–2511.

Humes LE, Busey TA, Craig JC, Kewley-Port D. The effects of age on sensory thresholds and temporal gap detection in hearing, vision, and touch. Attention, Perception, & Psychophysics. 2009; 71:860–71.

Humes LE, Dubno JR. Factors affecting speech understanding in older adults. In: The Aging Auditory System. Springer, New York, NY. 2010; p. 211–257.

Humes LE, Kewley-Port D, Fogerty D, Kinney D. Measures of hearing threshold and temporal processing across the adult lifespan. Hearing Research. 2010; 264:30–40.

Hunter LL, Sanford CA. Tympanometry and Wideband Acoustic Immitance. In: Handbook of Clinical Audiology, 7th edition. 2015; p.137–163.

Kale S, Heinz MG. Envelope coding in auditory nerve fibers following noise-induced hearing loss. Journal of the Association for Research in Otolaryngology. 2010; 11:657–673.

Kong YY, Mullangi A, Ding N. Differential modulation of auditory responses to attended and unattended speech in different listening conditions. Hearing research. 2014; 316:73–81.

Lakatos P, Chen CM, O’Connell MN, Mills A, Schroeder CE. Neuronal oscillations and multisensory interaction in primary auditory cortex. Neuron. 2007; 53:279–92.

Lehmann A, Schönwiesner M. Selective attention modulates human auditory brainstem responses: relative contributions of frequency and spatial cues. PLoS One. 2014; 9:e85442

Lin FR, Yaffe K, Xia J, Xue QL, Harris TB, Purchase-Helzner E, Satterfield S, Ayonayon HN, Ferrucci L, Simonsick EM, Health ABC Study Group F. Hearing loss and cognitive decline in older adults. JAMA Internal Medicine. 2013; 173:293–299.

Luo H, Poeppel D. Phase patterns of neuronal responses reliably discriminate speech in human auditory cortex. Neuron. 2007; 54:1001–1010.

Mai G, Tuomainen J, Howell P. Relationship between speech-evoked neural responses and perception of speech in noise in older adults. The Journal of the Acoustical Society of America. 2018; 143:1333–1345.

Mai G, Schoof T, Howell P. Modulation of phase-locked neural responses to speech during different arousal states is age-dependent. NeuroImage. 2019; 189:734–744.

Martin N, Lafortune M, Godbout J, Barakat M, Robillard R, Poirier G, Bastien C, Carrier J. Topography of age-related changes in sleep spindles. Neurobiology of aging. 2013; 34:468–476.

Morillon B, Liégeois-Chauvel C, Arnal LH, Bénar CG, Giraud AL. Asymmetric function of theta and gamma activity in syllable processing: an intra-cortical study. Frontiers in Psychology. 2012; 3:248.

Noguchi Y, Fujiwara M, Hamano S. Temporal evolution of neural activity underlying auditory discrimination of frequency increase and decrease. Brain Topography. 2015; 28:437–444.

Ng BS, Logothetis NK, Kayser C. EEG phase patterns reflect the selectivity of neural firing. Cerebral Cortex. 2012; 23:389–398.

Oya H, Gander PE, Petkov CI, Adolphs R, Nourski KV, Kawasaki H, Howard MA, Griffiths TD. Neural phase locking predicts BOLD response in human auditory cortex. NeuroImage. 2018; 169:286–301.

Peelle JE, Gross J, Davis MH. Phase-locked responses to speech in human auditory cortex are enhanced during comprehension. Cerebral Cortex. 2013; 23:1378–1387.

Petersen RC, Smith GE, Waring SC, Ivnik RJ, Tangalos EG, Kokmen E. Mild cognitive impairment: clinical characterization and outcome. Archives of Neurology. 1999; 56(3):303–8.

Plomp R, Mimpen AM. Speech-reception threshold for sentences as a function of age and noise level. The Journal of the Acoustical Society of America. 1979; 66:1333–1342.

Portas CM, Krakow K, Allen P, Josephs O, Armony JL, Frith CD. Auditory processing across the sleep-wake cycle: simultaneous EEG and fMRI monitoring in humans. Neuron. 2000; 28:991–999.

Presacco A, Simon JZ, Anderson S. Evidence of degraded representation of speech in noise, in the aging midbrain and cortex. Journal of Neurophysiology. 2016; 116:2346–2355.

Presacco A, Simon JZ, Anderson S. Effect of informational content of noise on speech representation in the aging midbrain and cortex. Journal of Neurophysiology. 2016; 116:2356–2367.

Presacco A, Simon JZ, Anderson S. Speech-in-noise representation in the aging midbrain and cortex: Effects of hearing loss. PloS One. 2019; 14(3):e0213899.

Rimmele JM, Golumbic EZ, Schröger E, Poeppel D. The effects of selective attention and speech acoustics on neural speech-tracking in a multi-talker scene. Cortex. 2015; 68:144–154.

Schneider BA, Li L, Daneman M. How competing speech interferes with speech comprehension in everyday listening situations. Journal of the American Academy of Audiology. 2007; 18(7):559–72.

Schneider BA, Hamstra SJ. Gap detection thresholds as a function of tonal duration for younger and older listeners. The Journal of the Acoustical Society of America. 1999; 106:371–380.

Schoof T, Rosen S. The role of auditory and cognitive factors in understanding speech in noise by normal-hearing older listeners. Frontiers in Aging Neuroscience. 2014; 6:307.

Schoof T, Rosen S. The role of age-related declines in subcortical auditory processing in speech perception in noise. Journal of the Association for Research in Otolaryngology. 2016; 17:441–460.

Shinn-Cunningham BG, Best V. Selective attention in normal and impaired hearing. Trends in Amplification. 2008; 12(4):283–99.

Skoe E, Kraus N. Auditory brainstem response to complex sounds: a tutorial. Ear and Hearing. 2010; 31:302.

Sommers MS, Gehr SE. Auditory suppression and frequency selectivity in older and younger adults. The Journal of the Acoustical Society of America. 1998; 103:1067–74.

Stine RA. Graphical interpretation of variance inflation factors. The American Statistician. 1995; 49:53–56.

Thatcher RW. Coherence, phase differences, phase shift, and phase lock in EEG/ERP analyses. Developmental neuropsychology. 2012; 37:476–96.

Tlumak AI, Durrant JD, Delgado RE. The effect of advancing age on auditory middle-and long-latency evoked potentials using a steady-state-response approach. American journal of audiology. 2015; 24:494–507.

Tun PA, O’kane G, Wingfield A. Distraction by competing speech in young and older adult listeners. Psychology and Aging. 2002; 17:453.

Tun PA, McCoy S, Wingfield A. Aging, hearing acuity, and the attentional costs of effortful listening. Psychology and Aging. 2009; 24(3):761.

Waschke L, Wöstmann M, Obleser J. States and traits of neural irregularity in the age-varying human brain. Scientific reports. 2017; 7:17381.

Wilf M, Ramot M, Furman-Haran E, Arzi A, Levkovitz Y, Malach R. Diminished auditory responses during NREM sleep correlate with the hierarchy of language processing. PloS One. 2016; 11:e0157143.

Wong PC, Skoe E, Russo NM, Dees T, Kraus N. Musical experience shapes human brainstem encoding of linguistic pitch patterns. Nature Neuroscience. 2007; 10:420.

Zhong Z, Henry KS, Heinz MG. Sensorineural hearing loss amplifies neural coding of envelope information in the central auditory system of chinchillas. Hearing Research. 2014; 309:55–62.

