## Appendix 1 ANCOVA results and Appendix 2 Simulations that test relative reliability of FFR measurements for "Relationship between neural phase-locked responses to speech and perception of speech in noise in young and older adults"

ANCOVAs were conducted (with Greenhouse-Geisser correction) for the five EEG signatures ($FFR_{ENV\_F0}$, $FFR_{PLV\_F0}$, $FFR_{TFS\_H2}$, $FFR_{TFS\_F2F3}$, and Logit-θ-PLV), three Noise Types (Quiet, SpN and BbN) and Age Group (young vs. older) as factors, and PTA ($PTA_{Low}$, $PTA_{High}$ and $PTA_{Wide}$) as covariates. **Tables A1**, **A2** and **A3** summarize the statistics using $PTA_{Low}$, $PTA_{High}$ and $PTA_{Wide}$ as the covariate, respectively.

**Table A1** Statistical results of ANCOVAs with $PTA_{Low}$ as the covariate. DV, df, F, and *p* refer to the dependent variables, degrees of freedom, F values, and *p* values, respectively. Significant *p* values (< 0.05) are indicated in bold. * = $p < 0.05$; ** = $p < 0.01$; *** = $p < 0.001$.

| DV | Factors | *df1* | *df2* | *F* | *p* |
|---|---|---|---|---|---|
| $FFR_{ENV\_F0}$ | Noise Type | 1.506 | 39.152 | 50.277 | **< $10^{-9}$***** |
| | Age Group | 1 | 26 | 8.842 | **0.006**** |
| | $PTA_{Low}$ | 1 | 26 | 12.943 | **0.001**** |
| | Noise Type × Age Group | 1.506 | 39.152 | 3.118 | 0.069 |
| | Noise Type × $PTA_{Low}$ | 1.506 | 39.152 | 1.445 | 0.246 |
| $FFR_{PLV\_F0}$ | Noise Type | 1.494 | 38.836 | 37.842 | **< $10^{-7}$***** |
| | Age Group | 1 | 26 | 1.196 | 0.284 |
| | $PTA_{Low}$ | 1 | 26 | 0.005 | 0.943 |
| | Noise Type × Age Group | 1.494 | 38.836 | 1.290 | 0.279 |
| | Noise Type × $PTA_{Low}$ | 1.494 | 38.836 | 0.372 | 0.630 |
| $FFR_{TFS\_H2}$ | Noise Type | 1.941 | 50.463 | 0.477 | 0.618 |

| | | | | | |
|---|---|---|---|---|---|
| | Age Group | 1 | 26 | 0.041 | 0.840 |
| | $PTA_{Low}$ | 1 | 26 | 0.050 | 0.824 |
| | Noise Type × Age Group | 1.941 | 50.463 | 2.698 | 0.079 |
| | Noise Type × $PTA_{Low}$ | 1.941 | 50.463 | 2.839 | 0.069 |
| $FFR_{TFS\_F2F3}$ | Noise Type | 1.487 | 38.654 | 1.493 | 0.237 |
| | Age Group | 1 | 26 | 2.035 | 0.166 |
| | $PTA_{Low}$ | 1 | 26 | 0.192 | 0.665 |
| | Noise Type × Age Group | 1.487 | 38.654 | 0.626 | 0.516 |
| | Noise Type × $PTA_{Low}$ | 1.487 | 38.654 | 0.123 | 0.855 |
| Logit-θ-PLV | Noise Type | 1.720 | 44.722 | 29.828 | **< $10^{-7}$***** |
| | Age Group | 1 | 26 | 4.608 | **0.041*** |
| | $PTA_{Low}$ | 1 | 26 | 1.390 | 0.249 |
| | Noise Type × Age Group | 1.720 | 44.722 | 0.267 | 0.610 |
| | Noise Type × $PTA_{Low}$ | 1.720 | 44.722 | 0.382 | 0.542 |

**Table A2** Statistical results of ANCOVAs with $PTA_{High}$ as the covariate. DV, df, F, and *p* refer to the dependent variables, degrees of freedom, F values, and *p* values, respectively. Significant *p* values (< 0.05) are indicated in bold. * = $p < 0.05$; ** = $p < 0.01$; *** = $p < 0.001$.

| DV | Factors | *df1* | *df2* | *F* | *p* |
|---|---|---|---|---|---|
| $FFR_{ENV\_F0}$ | Noise Type | 1.550 | 40.290 | 52.028 | **< $10^{-12}$***** |
| | Age Group | 1 | 26 | < 0.001 | 0.978 |
| | $PTA_{High}$ | 1 | 26 | 0.168 | 0.670 |
| | Noise Type × Age Group | 1.550 | 40.290 | 0.534 | 0.551 |
| | Noise Type × $PTA_{High}$ | 1.550 | 40.290 | 1.704 | 0.199 |
| $FFR_{PLV\_F0}$ | Noise Type | 1.517 | 39.444 | 37.385 | **< $10^{-7}$***** |

| | | | | | |
|---|---|---|---|---|---|
| | Age Group | 1 | 26 | 0.455 | 0.506 |
| | $PTA_{High}$ | 1 | 26 | 0.190 | 0.666 |
| | Noise Type × Age Group | 1.517 | 39.444 | 0.900 | 0.389 |
| | Noise Type × $PTA_{High}$ | 1.517 | 39.444 | 0.161 | 0.792 |
| $FFR_{TFS\_H2}$ | Noise Type | 1.902 | 49.465 | 0.381 | 0.675 |
| | Age Group | 1 | 26 | 0.247 | 0.623 |
| | $PTA_{High}$ | 1 | 26 | 0.384 | 0.560 |
| | Noise Type × Age Group | 1.902 | 49.465 | 1.852 | 0.169 |
| | Noise Type × $PTA_{High}$ | 1.902 | 49.465 | 0.492 | 0.605 |
| $FFR_{TFS\_F2F3}$ | Noise Type | 1.600 | 41.594 | 1.564 | 0.219 |
| | Age Group | 1 | 26 | 2.678 | 0.114 |
| | $PTA_{High}$ | 1 | 26 | 1.019 | 0.322 |
| | Noise Type × Age Group | 1.600 | 41.594 | 0.059 | 0.909 |
| | Noise Type × $PTA_{High}$ | 1.600 | 41.594 | 0.030 | 0.946 |
| Logit-θ-PLV | Noise Type | 1.759 | 45.731 | 32.811 | **$< 10^{-8}$***** |
| | Age Group | 1 | 26 | 5.937 | **0.022*** |
| | $PTA_{High}$ | 1 | 26 | 0.017 | 0.897 |
| | Noise Type × Age Group | 1.759 | 45.731 | 0.267 | 0.610 |
| | Noise Type × $PTA_{High}$ | 1.759 | 45.731 | 2.922 | 0.070 |

**Table A3** Statistical results of ANCOVAs with $PTA_{Wide}$ as the covariate. DV, df, F, and *p* refer to the dependent variables, degrees of freedom, F values, and *p* values, respectively. Significant *p* values (< 0.05) are indicated in bold. * = *p* < 0.05; ** = *p* < 0.01; *** = *p* < 0.001.

| DV | Factors | *df1* | *df2* | *F* | *p* |
|---|---|---|---|---|---|
| $FFR_{ENV\_F0}$ | Noise Type | 1.487 | 38.674 | 52.205 | **$< 10^{-12}$***** |

| | | | | | |
|---|---|---|---|---|---|
| | Age Group | 1 | 26 | 1.278 | 0.269 |
| | $PTA_{Wide}$ | 1 | 26 | 0.967 | 0.334 |
| | Noise Type × Age Group | 1.487 | 38.674 | 0.899 | 0.388 |
| | Noise Type × $PTA_{Wide}$ | 1.487 | 38.674 | 1.960 | 0.164 |
| $FFR_{PLV\_F0}$ | Noise Type | 1.523 | 39.602 | 37.268 | **$< 10^{-7}$***** |
| | Age Group | 1 | 26 | 0.405 | 0.530 |
| | $PTA_{Wide}$ | 1 | 26 | 0.139 | 0.713 |
| | Noise Type × Age Group | 1.523 | 39.602 | 0.610 | 0.505 |
| | Noise Type × $PTA_{Wide}$ | 1.523 | 39.602 | 0.070 | 0.887 |
| $FFR_{TFS\_H2}$ | Noise Type | 1.930 | 50.182 | 0.488 | 0.617 |
| | Age Group | 1 | 26 | 0.242 | 0.627 |
| | $PTA_{Wide}$ | 1 | 26 | 0.315 | 0.580 |
| | Noise Type × Age Group | 1.930 | 50.182 | 2.313 | 0.111 |
| | Noise Type × $PTA_{Wide}$ | 1.930 | 50.182 | 1.421 | 0.251 |
| $FFR_{TFS\_F2F3}$ | Noise Type | 1.581 | 41.114 | 1.450 | 0.245 |
| | Age Group | 1 | 26 | 2.777 | 0.108 |
| | $PTA_{Wide}$ | 1 | 26 | 0.970 | 0.334 |
| | Noise Type × Age Group | 1.581 | 41.114 | 0.080 | 0.883 |
| | Noise Type × $PTA_{Wide}$ | 1.581 | 41.114 | 0.265 | 0.716 |
| Logit-θ-PLV | Noise Type | 1.765 | 45.902 | 30.808 | **$< 10^{-7}$***** |
| | Age Group | 1 | 26 | 3.516 | 0.072 |
| | $PTA_{Wide}$ | 1 | 26 | 0.377 | 0.545 |
| | Noise Type × Age Group | 1.765 | 45.902 | 0.375 | 0.663 |
| | Noise Type × $PTA_{Wide}$ | 1.765 | 45.902 | 2.236 | 0.124 |

### Appendix 2. Simulations that test relative reliability of FFR measurements

Simulations were conducted to test which measurements – the FFR magnitudes, or response SNRs (difference in magnitudes between FFRs and the EEG noise floors) – can more reliably quantify FFRs.

#### Appendix 2.1 Methods

Simulated FFRs and EEG background noise were created. FFRs were created based on the stimulus syllable /i/ used in the present study: $FFR_{ENV}$ used the Hilbert Envelope of the syllable, while $FFR_{TFS}$ used the syllable *per se*. EEG background noise was pink noise (with random phases) that fits the 1/f power law of EEG. This artificially created 'real' FFRs (FFRs before adding EEG noise) and 'observed' FFRs (FFRs after adding EEG noise). The 'observed' FFR magnitudes and 'observed' SNRs (difference in magnitudes between 'observed' FFRs and EEG noise floors) were then measured in order to investigate which one can better reflect the 'real' FFR magnitudes. Measurements of the 'real' and 'observed' FFR magnitudes followed the same procedure as described in *2.4.1* in the main text. EEG noise floors were measured as EEG noise magnitudes at the corresponding frequency range of FFRs (110 ~ 160 Hz ($F_0$) for $FFR_{ENV\_F0}$; 220 ~ 320 Hz (H2) for $FFR_{TFS\_H2}$; 150-Hz bandwidth centred at 2400 Hz (F2) and 300-Hz bandwidth centred at 3100 Hz (F3) for $FFR_{TFS\_F2F3}$) at the 50-ms FFR pre-stimulus period (see *2.4.5* in the main text; also see Schoof & Rosen, 2016; Mai et al., 2018).

100 pairs of FFRs and EEG noise with different magnitudes were created and were made sure that they were in line with the characteristics of actual data in the present experiment: (i) the 'observed' FFRs had significantly greater magnitudes than the noise floors (see *2.4.5*); (ii) average magnitudes of the 'observed' FFRs and the noise floors approximated those of the actual data in the experiment (averaged across all participants) (see **Table A4**). The 'real' FFR magnitudes were then correlated with the 'observed' FFR magnitudes and 'observed' SNRs, where higher correlation values indicate better reliability of measurements. Such simulations were repeated 20 times and pair-wise t-tests were finally conducted to compare the two correlations (correlations between the 'real' FFR magnitudes and 'observed' FFR magnitudes vs. correlations between the 'real' FFR magnitudes and 'observed' SNRs; Fisher-transformed) for each of the three FFR signatures ($FFR_{ENV\_F0}$, $FFR_{TFS\_H2}$ and $FFR_{TFS\_F2F3}$) under each noise type (Quiet, SpN and BbN).

**Table A4.** Average magnitudes of the observed FFRs, EEG noise floors and SNRs of the actual and simulated data (numbers in the brackets indicate the simulated data) for the three FFR signatures ($FFR_{ENV\_F0}$, $FFR_{TFS\_H2}$ and $FFR_{TFS\_F2F3}$) under different noise types (Quiet, SpN and BbN). As shown, the average simulated magnitudes of the 'observed' FFRs, noise floors and SNRs approximate those of the actual data in the experiment.

| FFR signatures | Noise types | Observed FFR magnitudes | Magnitudes of EEG noise floors | SNRs |
|---|---|---|---|---|
| $FFR_{ENV\_F0}$ | Quiet | 4.771 (4.770) | 3.022 (3.041) | 1.749 (1.729) |
| | SpN | 3.697 (3.679) | 3.062 (3.055) | 0.635 (0.624) |
| | BbN | 3.752 (3.762) | 3.222 (3.220) | 0.530 (0.542) |
| $FFR_{TFS\_H2}$ | Quiet | 4.164 (4.160) | 2.362 (2.327) | 1.802 (1.834) |
| | SpN | 4.170 (4.180) | 2.270 (2.313) | 1.901 (1.867) |
| | BbN | 4.127 (4.132) | 2.329 (2.353) | 1.797 (1.779) |
| $FFR_{TFS\_F2F3}$ | Quiet | -30.632 (-30.694) | -37.884 (-37.841) | 7.252 (7.147) |
| | SpN | -30.341 (-30.512) | -38.192 (-38.143) | 7.852 (7.631) |
| | BbN | -30.389 (-30.313) | -37.730 (-37.784) | 7.341 (7.471) |

### Appendix 2.2 Results

Results showed that correlations between the 'real' FFR magnitudes and 'observed' FFR magnitudes were significantly higher than correlations between the 'real' FFR magnitudes and 'observed' SNRs for all three FFR signatures in all noise types ($FFR_{ENV\_F0\_Quiet}$, $p < 10^{-10}$; $FFR_{ENV\_F0\_SpN}$, $p < 10^{-5}$; $FFR_{ENV\_F0\_BbN}$, $p < 10^{-7}$; $FFR_{TFS\_H2\_Quiet}$, $p < 10^{-12}$; $FFR_{TFS\_H2\_SpN}$, $p < 10^{-9}$; $FFR_{TFS\_H2\_BbN}$, $p < 10^{-10}$; $FFR_{TFS\_F2F3\_Quiet}$, $p < 10^{-11}$; $FFR_{TFS\_F2F3\_SpN}$, $p < 10^{-15}$; $FFR_{TFS\_F2F3\_BbN}$, $p < 10^{-13}$) (**Figure A1**). This therefore indicates that, compared to the response SNR, the observed FFR magnitude should more reliably quantify the real FFR magnitude in the present study.

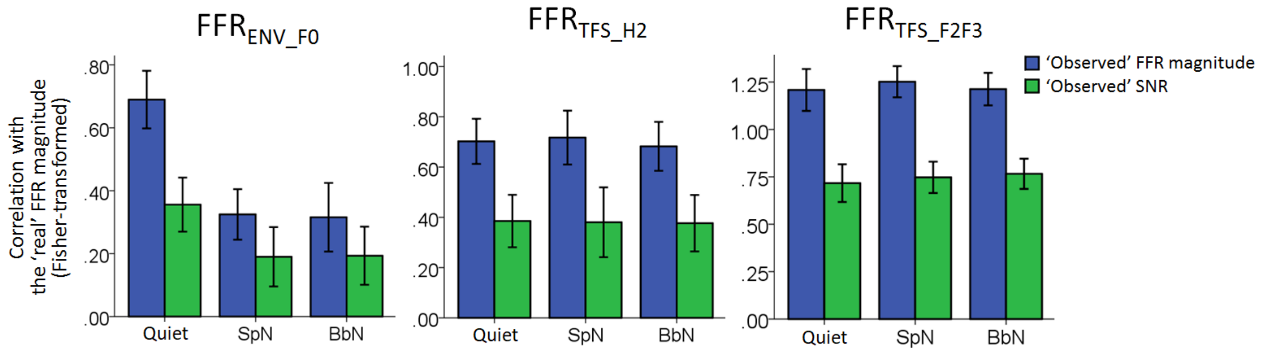


**Figure A1. Relative reliability of FFR measurements.** The 'real' FFR magnitudes were correlated (Fisher-transformed) with the 'observed' FFR magnitudes (*blue*) and the 'observed' SNRs (*green*) for the three FFR signatures ($FFR_{ENV\_F0}$, $FFR_{TFS\_H2}$ and $FFR_{TFS\_F2F3}$) under each noise type (Quiet, SpN and BbN). Error bars denote the standard deviations across the 20 simulations.
